# A platform for the recombinant production of Group A Streptococcus glycoconjugate vaccines

**DOI:** 10.1101/2024.03.01.582896

**Authors:** Sowmya Ajay Castro, Ian J. Passmore, Didier Ndeh, Helen Alexandra Shaw, Alessandro Ruda, Keira Burns, Sarah Thomson, Rupa Nagar, Kathirvel Alagesan, Kieron Lucas, Sherif Abouelhadid, Mark Reglinski, Ulrich Schwarz-Linek, Fatme Mawas, Göran Widmalm, Brendan W. Wren, Helge C. Dorfmueller

**Affiliations:** Division of Molecular Microbiology, School of Life Sciences, Dow Street, Dundee, DD1 5EH, United Kingdom; Department of Infection Biology, London School of Hygiene and Tropical Medicine, London, United Kingdom; The Medicines and Healthcare products Regulatory Agency (MHRA), Vaccines Division, Scientific Research & Innovation Group, Blanche Lane, South Mimms, Potters Bar, London EN6 3QG, United Kingdom; Department of Organic Chemistry, Stockholm University, SE-106 91 Stockholm, Sweden; Biological Services, School of Life Sciences, University of Dundee, DD1 5EH Dundee, United Kingdom; Max Planck Unit for the Science of Pathogens, Charitéplatz 1, 10117 Berlin, Germany; Biomedical Sciences Research Complex, University of St. Andrews, North Haugh, St. Andrews, Fife KY16 9ST, United Kingdom

## Abstract

Strep A is a human-exclusive bacterial pathogen killing annually more than 500,000 patients, and no current licensed vaccine exists. Strep A bacteria are highly diverse, but all produce an essential, abundant, and conserved surface carbohydrate, the Group A Carbohydrate, which contains a rhamnose polysaccharide (RhaPS) backbone. RhaPS is a validated universal vaccine candidate in a glycoconjugate prepared by chemical conjugation of the native carbohydrate to a carrier protein. We engineered the Group A Carbohydratte biosynthesis pathway to enable recombinant production using the industry standard route to couple RhaPS to selected carrier proteins within *E. coli* cells. The structural integrity of the produced recombinant glycoconjugate vaccines was confirmed by NMR spectroscopy and mass spectrometry. Purified RhaPS glycoconjugates elicited carbohydrate-specific antibodies in mice and rabbits and bound to the surface of multiple Strep A strains of diverse M-types, confirming the recombinantly produced RhaPS glycoconjugates as valuable vaccine candidates.

## INTRODUCTION

*Streptococcus pyogenes,* also known as Group A Streptococcus (Strep A or GAS), is associated with numerous diseases in humans ranging from more than 500 million cases per annum of pharyngitis to life-threatening invasive infections, such as streptococcal toxic syndrome and necrotizing fasciitis [1]. It is the primary cause of three post-infection autoimmune diseases; glomerulonephritis, rheumatic fever, and rheumatic heart disease. Strep A bacteria are responsible for over 500,000 deaths globally each year, particularly in low-income countries [1]. Antimicrobial resistance is emerging globally, and a Strep A vaccine is a current global imperative that the WHO has made a vaccine priority [2]. Hesitation in developing Strep A specific vaccines has been partly due to potential immune complications of vaccine components that may indirectly induce autoimmune complications and potentially lead to rheumatic heart disease [3]. Furthermore, an effective Strep A vaccine ideally needs to cover all >150 known variant serotypes (M-types) and be low-cost to be deployed in low resource countries where it is most needed.

All Strep A serotypes display the Lancefield Group A Carbohydrate (GAC) on the surface, which is conserved and immunogenic and therefore an excellent vaccine candidate [4–10]. GAC is a peptidoglycan-anchored surface structure consisting of a polyrhamnose backbone with *N*-acetylglucosamine (GlcNAc) side chains (Fig.1)[11, 12]. GAC biosynthesis is conducted in two discrete steps: In the cytosol, GacBCFG synthesise the GAC polyrhamnose backbone on the lipid precursor Und-PP-GlcNAc, which is then flipped across the membrane via the GacDE ABC-transporter [6, 13]. Outside the cell, the GlcNAc side chains are introduced via GacL and are further modified occasionally with glycerol phosphate groups (Fig.1)[14], before being linked to peptidoglycan via an unknown ligase [12]. However, the immunogenic GlcNAc side chains may, in combination with other Strep A components, be responsible for the Strep A specific highly complex post infection autoimmune diseases such as rheumatic fever and rheumatic heart disease, which kill estimated more than 250,000 patients annually [15]. A key desire for Strep A vaccine development therefore is to exclude any epitopes that could potentially trigger antibodies that cross react with the human tissue.

**Fig. 1.**
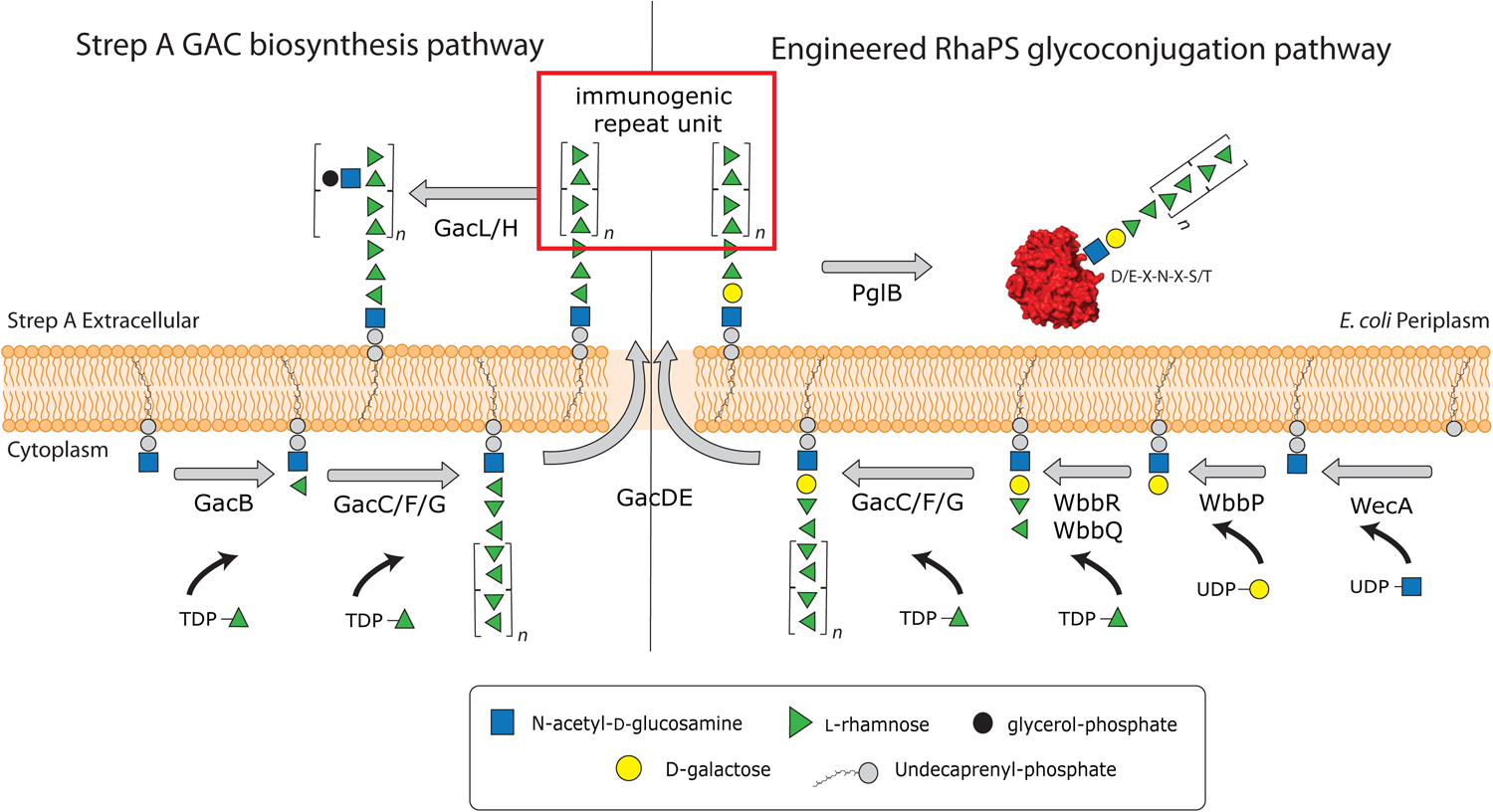
Schematic showing GAC biosynthesis pathway (left) and the revised GAC biosynthesis pathway (right) that synthesises OTase compatible reducing end sugars. The GAC pathway synthesises two major cell wall components, i) the undecorated immunogenic repeating unit (red box) and ii) the final GAC. Our engineered pathway includes the genes *wbbPRQ* from the *Shigella dysenteriae* serotype 1 O-antigen pathway. These glycosyltransferases introduce a different end group, α-L-Rha*p*-(1→3)-α-L-Rha*p*-(1→2)-α-D-Gal*p*-(1→3)-D-Glc*p*NAc that can be extended by GacCFG to produce the polyrhamnose backbone containing the immunogenic →2)-α-L-Rha*p*-(1→3)-α-L-Rha*p*-(1→ repeating unit (red box). Once the lipid linked product has been translocated across the cell membrane via GacDE (ABC-transporter), the OTase PglB recognises the reducing end group disaccharides, cleaves the glycan off the lipid carrier and transfers it onto the OTase sequon on the acceptor protein of choice. This completes the synthesis of the first recombinant Strep A glycoconjugate vaccine candidates.

Deletion of the *gacI* gene in Strep A bacteria results in a polyrhamnose homopolymer consisting of the α-(1→2)- and α-(1→3)-linked residues in disaccharide repeating units (RhaPS) that lacks the GlcNAc side chains and antibodies raised against this epitope do not trigger cross-reactivity with human tissue [6, 16]. RhaPS is found in Strep A and the closely related Group C streptococci [17] and has been previously validated as a Strep A vaccine candidate [18]. However, until now RhaPS vaccine candidates are produced using elaborate chemical conjugation methods or are chemically synthesised as up to two repeating units [4, 19, 20].

Glycoconjugate vaccines against bacteria are one of the success stories of modern medicine due to their ability to activate B- and (carbohydrate specific) T-cells [21], which can result in a long-lasting immune response. This has led to a significant reduction in the global occurrence of bacterial meningitis and pneumonia [22]. Historically, glycoconjugate vaccines are produced by covalently linking a bacterial polysaccharide to a carrier protein using complex multi-step chemical procedures that are expensive, laborious, and require several purification steps that could lead to batch-to-batch variation. Additionally, some chemical conjugation methods such as reductive amination can alter the polysaccharide epitopes, thereby affecting the immunogenicity of the glycoconjugate [23]. Only four glycoconjugate vaccines have been fully licenced to date, against *Haemophilus influenzae* type b, *Neisseria meningitidis*, *Streptococcus pneumoniae* and *Salmonella typhi* [24–26]. The continued use of the same carrier proteins (*e.g.,* the non-toxic mutant of diphtheria toxin, CRM197, and the inactivated tetanus toxoid, TT) in the burgeoning development of glycoconjugate vaccines has issues including pre-existing immunity to a given carrier protein which can counter the immune response to new glycoconjugate vaccines with the same carrier [27]. This carrier-induced epitopic suppression has been observed in children who exhibited a superior immune response when immunised with a CRM197-Hib glycoconjugate compared to those who received TT-Hib [28].

However, glycoconjugate vaccine design is set to change through the glycoengineering of recombinant vaccines in bacteria, mini factories to produce an inexhaustible and renewable supply of pure vaccine products [29]. This process is known as Protein Glycan Coupling Technology (PGCT), offering low-cost options to recombinantly produce customised glycoconjugates. Three required PGCT components are encoded in the bacteria for (i) the biosynthesis of the candidate glycan (usually capsular polysaccharide or O-antigen), (ii) candidate carrier protein containing sequon(s) to site-specific receive the glycan, and (iii) the coupling oligosaccharyltransferase (OTase) enzyme, for instance the first discovered OTase from *Campylobacter jejuni* (*Cj*PglB) [29–31]. Additional OTases have been reported from *Neisseria meningitidis* (PglL)[32] and more recently two OTases from *Acinetobacter* (PglL and PglS)[33–35]. PGCT could be applied to produce double-hit glycoconjugate vaccines where the glycan and carrier protein are from the same or dual-hit glycoconjugate vaccines where the glycan and carrier protein are from different pathogens, thus potentially providing protection from several serotypes or two pathogens. However, one drawback of PGCT is the relatively limited glycan substrate specificity of the OTase enzymes. For instance, *Cj*PglB exhibits poor transfer of glycans that lack an acetamido group modification on their reducing end sugar [30] or polysaccharides where the two sugars proximal to the lipid carrier are connected via a β-(1→4)-linkage [36]. Despite these limitations numerous vaccine candidates have now been engineered using PGCT, which include those improving existing licenced vaccines (*e.g.*, *Streptococcus pneumoniae*) [37, 38], entirely new vaccines for both Gram-positive [39] and Gram-negative bacteria [40–42] and (because of the low production costs) veterinary pathogens [43–45]. Several PGCT-derived vaccines are now in human clinical trials [46–48]. All previously reported PGCT produced vaccine candidates are built via the Wzx/Wzy-dependent pathway. These glycans are synthesised from oligosaccharides having 4-6 sugars, synthesised on the lipid carrier Und-PP in the cytosoland translocated across the lipid bilayer into the periplasm where they are extended by the Wzy-polymerase to form the final polysaccharide [49, 50].

In this study, we demonstrate production of a novel recombinant Group Strep A glycoconjugate vaccine by coupling the RhaPS glycan to Strep A-specific and non-Strep A carrier proteins using PGCT. We overcome the limitations of OTase (*Cj*PglB) reducing end group specificity by utilising glycosyltransferases from two *Shigella* O-antigen biosynthetic pathways to engineer two hybrid RhaPS glycans with distinct reducing end linkers: α-L-Rha*p*-(1→3)-α-L-Rha*p*-(1→2)-α-D-Gal*p*-(1→3)-β-D-Glc*p*NAc (*Shigella dysenteriae* type 1, Shi1) and α-L-Rha*p*-(1→3)-α-L-Rha*p*-(1→3)-β-D-Glc*p*NAc (*Shigella flexneri* type 2a, Shi2a). We demonstrate that these modified RhaPS have the requisite stereochemistry of the disaccharide reducing end group for coupling to carrier proteins by *Cj*PglB. We confirm the structural integrity of the recombinantly build RhaPS on the carrier proteins via NMR, mass spectrometry and immunoblotting and undertake initial immunological studies that reveal that these recombinantly synthesised glycoconjugates produce Strep A specific antibodies. Our platform allows the recombinant production of the first recombinant Strep A dual-hit vaccines and enables *via* its modular system the easy exchange of the carrier protein with other pathogen specific proteins.

## RESULTS

### Engineering of OTase-compatible reducing end group linkers onto RhaPS from Strep A

We have previously cloned the requisite genes for Strep A RhaPS and demonstrated its heterologous production in *E. coli* [13]. RhaPS is synthesised by the enzymes GacB-G on the inner membrane as lipid-linked carbohydrate [9]. Production of RhaPS in an O-antigen ligase (*waaL*) positive strain of *E. coli* results in its deposition on the cell surface, which can serve as a platform for production of Outer Membrane Vesicle (OMV) vaccines [9]. The native GAC structure contains a rhamnose residue that is β-(1→4)-linked to GlcNAc at its reducing end (Fig.1)[13], which we hypothesised would not constitute a substrate for *Cj*PglB. All attempts to construct a native RhaPS glycoconjugate using *Cj*PglB were unsuccessful (data not shown). Therefore, we sought to modify the reducing end of RhaPS to generate a structure that would serve as a substrate for *Cj*PglB.

The *S. dysenteriae* type 1 and *S. flexneri* 2a O-polysaccharides have previously been utilised to generate glycoconjugates by PGCT and have been validated as substrates for *Cj*PglB [49, 50]. These structures have been characterised and gene/enzyme functions assigned accordingly [49, 50]. Since both *Shigella* O-antigen structures contain two rhamnose units at their non-reducing end, we attempted to create hybrid glycans that combine the *Shigella* repeating unit with the RhaPS structure from GAC, with the overall aim of generating a RhaPS glycan that would serve as a substrate for *Cj*PglB (Figure 1).

Thus, we replaced *gacB* from the Strep A GAC biosynthetic pathway (Fig.1) [6, 13] with two partial pathways: i) *S. dysenteriae* 1 genes *wbbP, wbbR* and *wbbQ*, encoding the O-antigen glycosyltransferases responsible for synthesis of lipid-linked α-L-Rha*p*-(1→3)-α-L-Rha*p*-(1→2)-α-D-Gal*p*-(1→3)-α-D-Glc*p*NAc (denoted Shi1 hereafter)(Fig.1)[49]; ii) *S. flexneri* 2a enzymes RfbF and RfbG, responsible for synthesis of lipid-linked α-L-Rha*p*-(1→3)-α-L-Rha*p*-(1→3)-α-D-Glc*p*NAc (denoted Shi2a hereafter)[50].

We have previously shown that deletion of *gacB* abrogates production of RhaPS [13, 51]. We now assessed the ability of the Shi1 and Shi2a repeating unit pathways to complement this *gacB* deletion (Fig.2A). Transformation of *E. coli* with plasmids containing either the Shi1 or Shi2a pathways restored RhaPS production as determined by immunoblot analysis (Fig.2A). This suggests that the *Shigella* genes are able to replace the function of *gacB* towards the biosynthesis of lipid-linked GAC (Fig.1). Deletion of either *wbbP* or *wbbQ* resulted in no GAC production, suggesting that all three genes are required for assembly (Fig.2B).

**Fig. 2.**
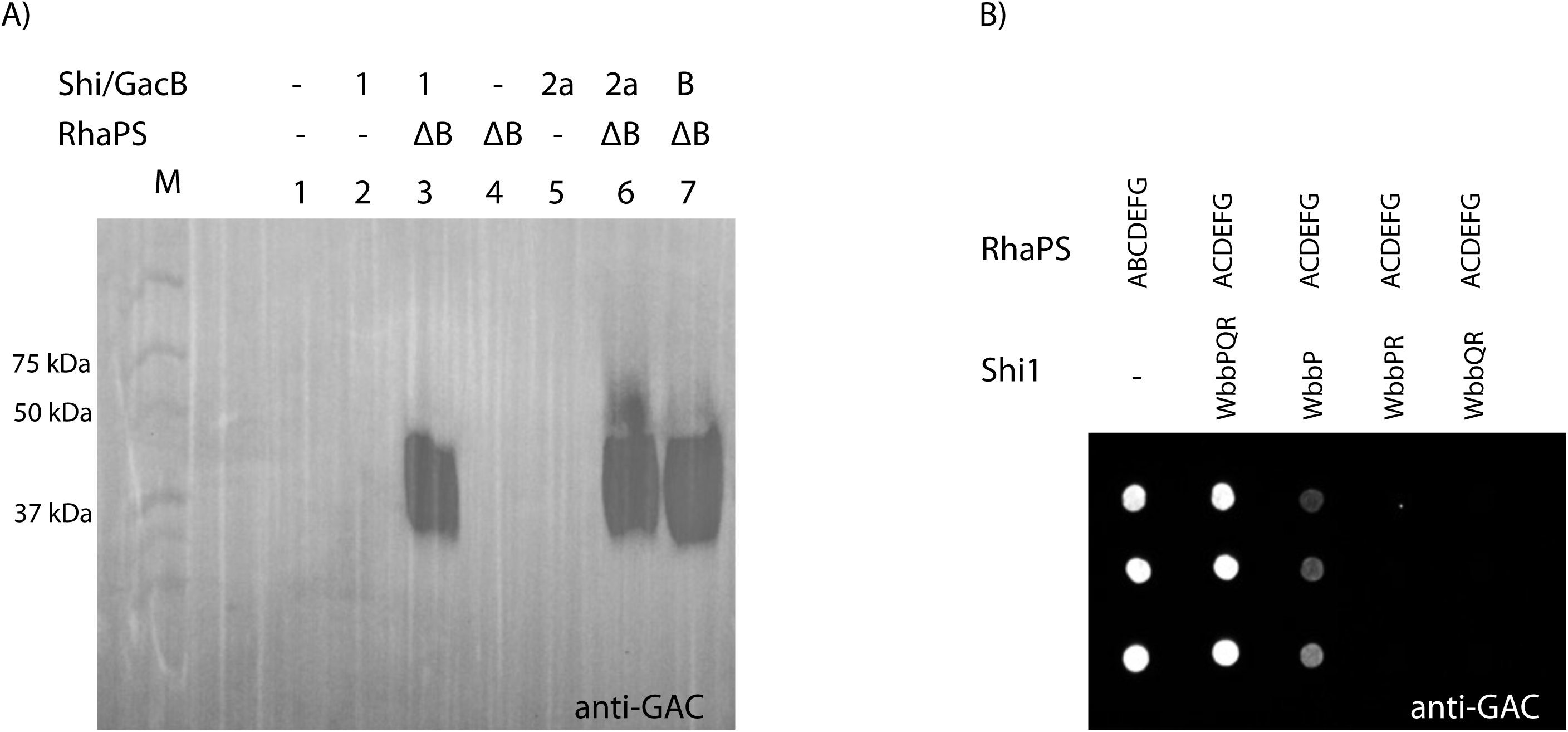
A) *E. coli rfaS* deletion cells transformed with two plasmids containing either the Shi1/2a gene cluster fragment, GacB control or empty plasmid (negative control) as well as the RhaPS clusters (ΔB) were grown overnight and whole cell lysates subjected to SDS-PAGE and western blotting with the commercially available anti-GAC antibodies. Plasmids indicated: ΔB = *gacACDEFG*; B = *gacB*; Shi1 = *wbbPQR*, Shi2a = *rfbFG*, and empty plasmid controls = -. B) *E. coli rfaS* deletion cells transformed with combinations of RhaPS cluster, and Shi1 gene cluster fragments were grown overnight and subjected to dot blot analysis using the GAC antibodies. RhaPS is synthesised by the WbbPQR proteins when co-expressed with the ΔB RhaPS proteins. Omitting one of the Shi1 proteins results in weaker or no RhaPS production.

Growth medium composition can influence glycan and glycoconjugate production [52]. Therefore, we explored recombinant glycan production in a range of standard growth media. Both, Shi1 and Shi2a built RhaPS were produced in all tested media (SupplFig.1A). Only the Shi2a-RhaPS glycan is built to a lesser extent in BHI media. We next aimed to determine whether Shi1/Shi2a engineered RhaPS would be recognised as substrates by *Cj*PglB. Plasmids encoding the RhaPS biosynthesis genes and the Shi1 or Shi2 repeating unit pathways were transformed into an *E. coli* strain that harboured a chromosomal, IPTG-inducible copy of *cjpglB* [53, 54].

Three candidate carrier proteins were selected as targets for coupling: *S. pneumoniae* neuraminidase, NanA [37], *S. pyogenes,* IdeS [55], and *P. aeruginosa* exotoxin, ExoA [39]. Each acceptor protein varied in its number of potential *N*-glycosylation sites. IdeS and NanA-constructs contain two *N-*glycosylation sequons [56] located on the N- and C-terminus, whilst ExoA contains two internal *N*-glycosylation sites and four N- and C-terminal sites (ExoA-10tag).

Each acceptor protein was tested in combination with both RhaPS-linkers and in a variety of growth media to determine optimal conditions for glycoconjugate production (Fig.3). Acceptor proteins were purified by His-affinity chromatography and were analysed by western blotting using protein-specific anti-His antibodies and glycan specific GAC antibodies (Fig.3; SupplFig.1B).

**Fig. 3.**
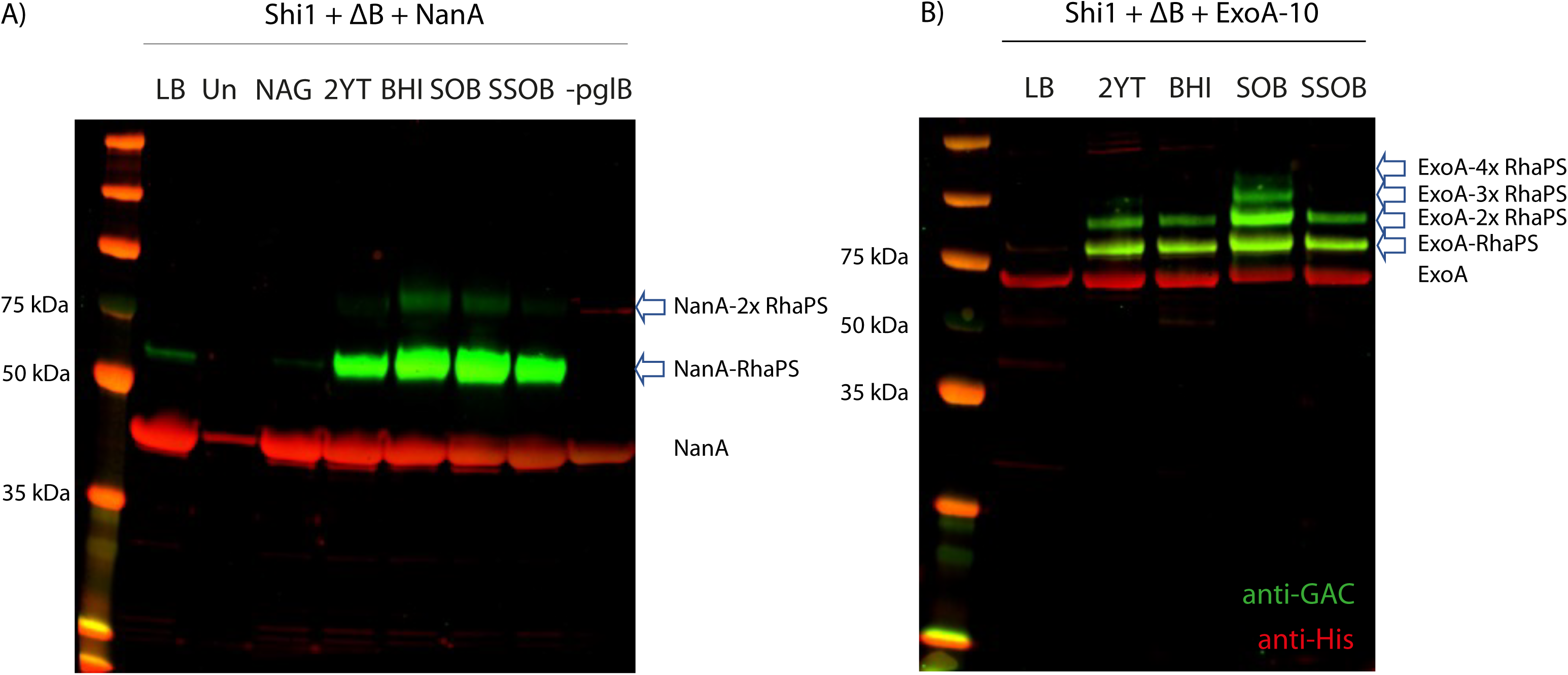
*E. coli* cells (Δ*waaL* with chromosomally integrated OTase *pglB* gene) were transformed with A) the Shi1 gene cluster, the RhaPS cluster (ΔB) and NanA carrier protein and B) Shi1, ΔB and ExoA-10 tag carrier protein plasmids. Cells were tested for glycoconjugate production in different growth conditions. Total cell lysates were run over SDS-PAGE and analysed via His and GAC-antibodies (red and green, respectively). Glycoconjugate production levels vary between the media conditions. Un = uninduced, -*pglB* = cells lack the chromosomal *pglB*. Arrows indicate glycosylated protein bands.

Glycosylation was observed for all three protein carriers (Fig.3; SupplFig.1B). This suggests that these hybrid RhaPS biosynthesis pathways generate glycan structures that are substrates for PglB. Optimum glycoconjugate yields varied for the three different carrier proteins and pathways (Fig.3; SupplFig.1B). NanA with the Shi1 pathway was glycosylated on both sites in SOB and BHI media, single site occupancy was observed in 2YT and SSOB media (Fig.3A). IdeS with the Shi2a pathway was mainly singly glycosylated (2YT, BHI, SOB media), whilst both sites were partially glycosylated with RhaPS in SSOB media (SupplFig1B). However, qualitatively lower glycoprotein yields were observed for IdeS-RhaPS compared with the other acceptor proteins-RhaPS combinations tested. ExoA shows a band pattern of higher molecular weight RhaPS-decorated species in SOB media, suggesting that four sites were glycosylated (Fig.3B). In 2YT, BHI and SSOB media two sites were glycosylated. The band-shifts between unglycosylated and glycosylated protein bands are estimated to reflect a 7-10 kDa increase in mass weight relative to the unglycosylated protein, consistent with the expected molecular weight of native RhaPS [11, 19](Fig.3). Taken together, these results suggest that reducing end modified RhaPS are viable substrates for *Cj*PglB and are compatible with current platforms for engineering of glycoconjugate vaccines in *E. coli*.

### Purification of IdeS-RhaPS and NanA-RhaPS glycoconjugates

The glycosylated proteins were purified for immunisation studies in mice and rabbits. In the first instance glycoconjugates were upscaled to 2 L cultures for NanA-RhaPS and IdeS-RhaPS for cobalt purification. NanA-RhaPS and IdeS-RhaPS fractions were analysed by SDS-PAGE and desalted into endotoxin-free PBS. The fractions contained a mixture of unglycosylated NanA-/IdeS and RhaPS-decorated proteins, with the majority being unglycosylated (SupplFig.2). These samples were used in immunisation studies to evaluate if the low yield of RhaPS-decorated proteins was sufficient to trigger (carbohydrate-specific) antibodies and whether the engineered glycoconjugates were able to stimulate long-lasting immunity. Affinity purified fractions containing NanA-RhaPS and IdeS-RhaPS, respectively, were pooled and purified by size exclusion chromatography to remove unglycosylated material and to obtain a greater proportion of RhaPS-conjugate protein. Fractions containing the NanA-RhaPS glycoconjugate eluted earlier than the unglycosylated carrier protein and were pooled for a second size exclusion chromatography (Fig.4A,B). Only those fractions containing the glycoconjugates were pooled and most unglycosylated carrier proteins were removed as judged by SDS-PAGE analysis (Fig.4A,B). The purification protocol for IdeS-RhaPS was also optimised, isolating the glycoconjugates with one and two glycosylation sites and removing excess unglycosylated protein (SupplFig.3). The concentrated NanA-RhaPS fractions were subsequently used for rabbit immunisation studies.

**Fig. 4.**
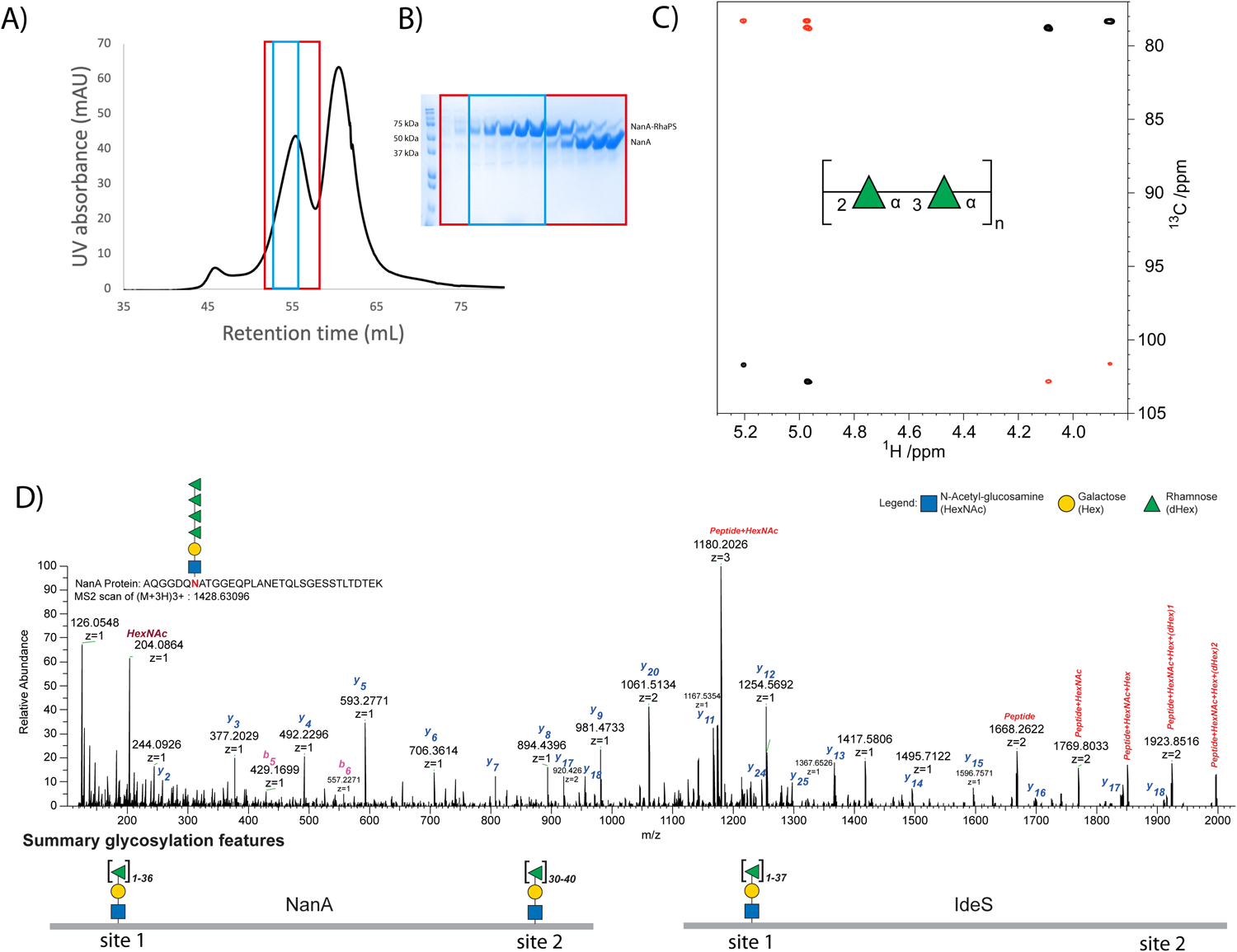
A) NTA column purified NanA-RhaPS was purified *via* two rounds of size exclusion chromatography to isolate the higher molecular weight glycosylated NanA-RhaPS species. B) Fractions of the second round were analysed via SDS-PAGE and reveal sequential elution of NanA-RhaPS glycoconjugate and free-protein species. The fractions boxed in blue were pooled for immunisation studies. C) NMR analysis con-firms that the glycoconjugate vaccine candidates carry the native backbone repeating unit with the structure [→2)-α-L-Rha*p*-(1→3)-α-L-Rha*p*-(1→]_n_. Selected region of overlaid ^1^H,^13^C-HSQC (black) and ^1^H,^13^C-HMBC (red) NMR spectra of the engineered glycoconjugate. The repeating unit of the rhamnan polysaccharide structure is given in SNFG format[71]. D) HCD spectra of a tryptic peptide derived from NanA protein. The monoisotopic mass of the precursor ion [M + 3H]^3+^ m/z 1428.63096 corresponds to the peptide AQGGDQNATGGEQPLANETQLSGESSTLTDTEK with Δmass 949 Da corresponding to HexNAc-Hex-(dHex)_4_. The lists of the b and y ions assigned from MS/MS spectra are shown in this figure. Summary glycosylation features indicate observed site-specific glycosylation for NanA and IdeS. We observed that the glycosylation site close to N-terminal for both proteins could carry up to 36 Rha residues. For the protein NanA, the second glycosylation site was mostly detected, carrying 30 to 40 Rha residues. The C-terminal glycosylation site of IdeS protein exhibited diverse glycan heterogeneity carrying 9 to 41 Rha residues.

### NMR spectroscopy confirms the structure of the polyrhamnose repeating unit

To ensure accurate reproduction of GAC when heterologously produced in *E. coli,* we conducted structural characterisation of the engineered glycoconjugates via mass spectrometry and NMR analysis. The carbohydrate composition analysis of the glycoconjugate by GC-MS showed the presence of rhamnose (SupplFig.4A) as the major carbohydrate component. The MS fragmentation pattern of the rhamnose derivative, eluting at RT 11.77 min, showed ions at *m/z* 204 and *m/z* 217 that are characteristic of the monosaccharide (SupplFig.4B)[57]. Subsequently, we conducted further analysis by NMR spectroscopy. ^1^H and ^13^C NMR chemical shifts of carbohydrate residues in polysaccharides on one hand and amino acids in polypeptides or proteins on the other show in many instances relatively limited spectral overlap. [58] Thus, it was possible to identify cross-peaks anticipated to derive from sugar residues. In the spectral region where resonances from anomeric atoms of sugar residues reside two cross-peaks were identified (Fig.4C), at δ_H_/δ_C_ 5.20/101.70 and δ_H_/δ_C_ 4.97/102.82, consistent with a rhamnose polysaccharide containing two sugar residues in the repeating unit. Furthermore, eight cross-peaks were observed in the spectral regions δ_H_ 4.2 – 3.5 and δ_C_ 79 − 70 arising from protons and carbons of two sugar rings. NMR resonances from the exocyclic methyl groups of rhamnosyl residues were not possible to unambiguously identify from the ^1^H,^13^C-HSQC spectrum due to the presence of cross-peaks originating from the protein. However, correlations were observed between δ_H_ 1.34 and δ_C_ 70.04 and 72.94, and between δ_H_ 1.29 and δ_C_ 70.10 and 72.40 in a ^1^H,^13^C-HMBC NMR spectrum, *i.e.*, consistent with protons of methyl groups in rhamnosyl residues and carbon atoms in the spectral region for sugars having a pyranoid ring form; consequently, the C6 NMR chemical shifts were possible to identify from the ^1^H,^13^C-HSQC NMR spectrum. In addition, the ^1^H,^13^C-HMBC NMR spectrum also revealed correlations between anomeric protons and glycosyloxylated carbons (∼78 ppm) as well as between anomeric carbons and protons at glycosyloxylated carbons (Fig.4C). The unassigned NMR chemical shift data from the ^1^H,^13^C-HSQC spectrum and those originating from anomeric atoms in the ^1^H,^13^C-HMBC NMR spectrum were given as input to the CASPER program [59, 60] together with two residues of L-rhamnopyranose, but without defining anomeric configuration or any linkage position. The result of the automated structural elucidation revealed →2)-α-L-Rha*p*-(1→3)-α-L-Rha*p*-(1→ as repeating unit structure (cf. CASPER report and NMR chemical shift data in Supplementary information table, SupplTable6). In this type of analysis, the top-ranked structural proposal is normalized to unity, *δ*_Rel_ = 1.0, and in the list of ‘normalized relative rankings’ a difference of at least ∼20% (*i.e.*, *δ*_Rel_ = 1.2) between the top-ranked structure and the second-best structure is required if the first is to be chosen as the correct one. The second ranked structure showed *δ*_Rel_ = 2.2, *i.e.*, well beyond the difference required to differentiate structural alternatives. Thus, 2D NMR spectroscopy experiments in conjunction with analysis using the CASPER program reveal a disaccharide repeating unit with alternating linkages of the rhamnosyl residues, identical to the reported structure of the polysaccharide backbone for the GAC[61] and in agreement to the western blot analysis (Fig.3,SupplFig.1B).

### Determination of RhaPS conjugation via mass spectrometry

Several native GAC samples have been isolated and characterised for their approximate total mass[11, 19]. We set out to investigate if the recombinantly produced RhaPS glycoconjugates are of the native size. The purified NanA-RhaPS and IdeS-RhaPS glycoconjugates were subjected to mass spectrometry to determine the sites of conjugation and glycan mass. Mass spectrometry methods can be used to determine the carbohydrate mass (and consequently the number of repeating units in the polysaccharide) of glycoconjugate vaccine candidates, as exemplified by Ravenscroft *et al.* [49, 50]. The Strep A RhaPS determined in SEC-MALS experiments to have an approximate average mass of 7.2 kDa [11, 19]. Considering that our *E. coli* system produced all required proteins for the RhaPS biosynthesis except the GacB enzyme (Fig.1), we hypothesised that the average number of repeating units of the glycan on each *N*-glycosylation site should be of comparable mass to the natively synthesised carbohydrate.

After proteolysis, NanA- and IdeS-RhaPS were analysed by LC-MS/MS. We employed an unbiased, MS-based methodology based on high-field asymmetric waveform ion mobility spectrometry (FAIMS) fractionation using varying compensation voltage (CV)(−20, −30, −35, −40, and −50) to increase the detection of modified peptides [62, 63]. The data were searched with Byonic using a wildcard setting restricted to asparagine to enable the identification of modified peptides. As expected, identified modified peptides exhibited Δmass of 146.0579 Da units indicating the presence of sequential deoxyhexose (rhamnose) (Fig.4D, SupplFig.5, SupplTable3). The masses of these glycan modifications agreed with the theoretical mass and the Shi1 reducing end group sugars. Interestingly, we observed that in higher repeating units of rhamnose in both samples NanA and IdeS, the glycopeptides had an additional mass of 56 Da. The MS/MS spectra of these identified peptides indicated that they were glycopeptides due to glycan fragmentation and the repeating pattern of 146.0579 Da difference. Of all FAIMS voltage tested, −35 V/-40V resulted in the highest unique glycopeptide identification rate. We identified 32 and 29 unique glycopeptides corresponding to NanA (AQGGDQN[Glycan]ATGGEQPLANETQLSGESSTLTDTEK) and IdeS (AQGGDQN[Glycan]ATGGIRY), respectively. Our data suggest that both glycoconjugates carry up to 37 rhamnose residues (SupplTable3), reflecting a total mass of up to 5.8 kDa (SupplFig.5). Under the same condition, we also identified glycosylated C-terminal peptide corresponding to the sequence REQGGDQNATGGHHHHHHHHHH for NanA protein. In the case of IdeS, the expected glycopeptides were detected as either NQTNGGDQNATGGHHHHHHHHHH or as TLSTGQDSWNQTNGGDQNATGGHHHHHHHHHH. Based on Δmass and glycan fragments, our data indicates that the peptide derived from NanA protein carries 30 to 40 rhamnose residues (SupplTable4), whereas the IdeS C-terminal glycopeptide carries 9 to 41 rhamnose residues (SupplTable5). We identified 22 and 23 unique C-terminal glycopeptides corresponding to NanA and IdeS, respectively. We also detected 11 unique glycopeptides within 1 Da mass tolerance of the expected glycan mass (SupplTable5). In summary, the herein applied FAIMS-assisted open search glycopeptide analysis unambiguously revealed that the two carrier proteins are decorated with the expected (Rha)_n_-Gal-GlcNAc glycan moiety, thus confirming that the Shi1 pathway builds a reducing end linker that could be further elaborated to produce a native-like RhaPS glycan (Fig.1, SupplFig.5).

**Fig. 5.**
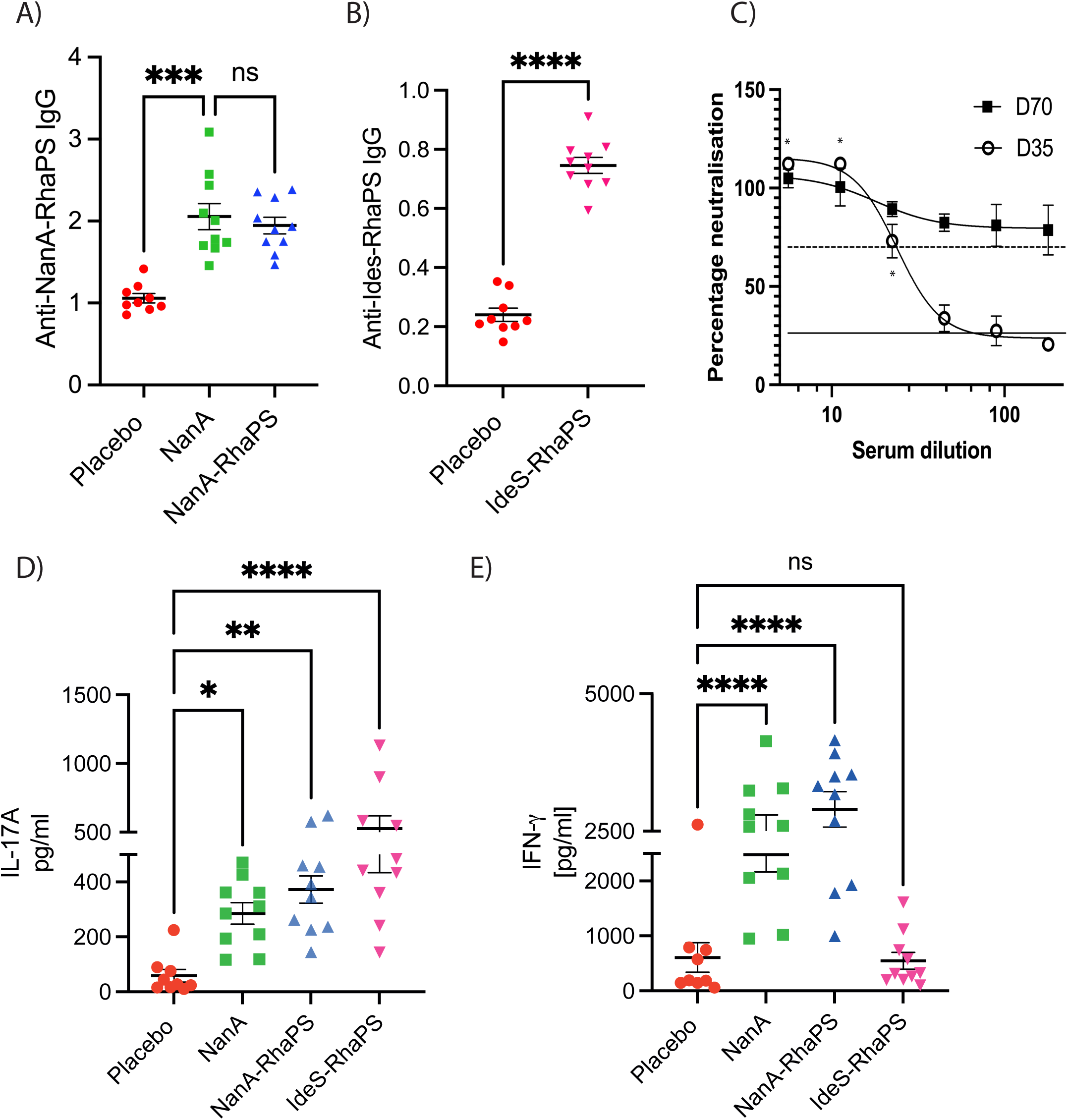
A,B) Mouse immunisation study (n = 10) (70 days). Mouse sera were investigated for IgG antibodies triggered post immunisation with the low RhaPS-containing NanA-RhaPS, NanA alone and with low RhaPS containing IdeS-RhaPS. Evaluation of vaccine candidate specific antibodies in mice sera using an ELISA based assay with respective glycoconjugate vaccine candidates showed that IgG levels are elevated in all immunized mice. Statistical analysis was performed using ANOVA followed by a Dunn’s post hoc test. *p<0.01; **P<0.001; ****P<0.0001; ns-not significant. C) Pooled mice sera (n = 10) from IdeS-RhaPS immunisation studies were tested in an IdeS neutralisation assay. A series of dilutions from 1:6 to 1:180 was tested in triplicate and the neutralisation activity is displayed in % compared with a no-sera control. * indicates duplicate data obtained for D35. Dashed line, positive control sera; solid line, negative control sera. D,E) Immunised mice splenocytes were investigated for IL17A and IFN-γ levels after restimulation with the mitogens NanA-RhaPS. Statistical analysis was performed using ANOVA followed by a Bonferroni’s post hoc test. *P<0.05; **P<0.001; ****P<0.0001.

### Evaluation of RhaPS glycoconjugates for their immunogenicity

We conducted mice immunisation studies (n=10) with three different vaccine candidates, NanA, NanA-RhaPS and IdeS-RhaPS. Our glycoconjugate immunisation studies showed that the vaccine candidates were well tolerated when administered in mice with three doses at days 0, 14 and 28. Post immunisation (Day 70), sera were analysed for immunogen specific antibodies (Fig.5A,B). All three candidates NanA, NanA-RhaPS and IdeS-RhaPS triggered antibodies significantly above the placebo vaccination levels, when evaluated in an ELISA assay. Considering that the carrier protein IdeS is a Strep A specific protein, we tested if IdeS-RhaPS also triggered IdeS specific antibodies that neutralise the protease activity of the enzyme [64]. All tested dilutions from Day 70 bleed showed that the IdeS protease activity was neutralised to >70% (Fig.5C) and exceeded the neutralisation ability of a positive control sample from previous IdeS immunisation [64]. The data suggests that IdeS-RhaPS could provide a dual-hit vaccine candidate, targeting the Strep A surface carbohydrate and the secreted protease IdeS.

In addition to the antibody profiling, we determined whether these conjugates were able to induce an immune cell response necessary for B cell activation ([21, 65]). We measured two markers of T cell activation (IFN-γ and IL-17) in isolated and restimulated splenocytes from the immunised mice. The data showed that significant levels of IL17A were triggered in animals immunised with NanA, NanA-RhaPS and IdeS-RhaPS, upon stimulation with the respective immunogen when compared to Mock (PBS) controls (Fig.5D,E). Moreover, the IFN-γ marker were shown to be significantly higher in all animals that received either NanA or NanA-RhaPS when compared to Mock controls (Fig5E). IdeS-RhaPS immunisation did not result in elevated IFN-γ levels, when compared to Mock controls. Therefore, an increase in both tested markers was noticed in spleen cells derived from animals that were immunised with NanA-RhaPS and NanA. Contrary, IdeS-RhaPS derived cells showed an increased level only in the IL17A cytokine mediators, suggesting a possible role for IL17A mediated immune response post immunisation (Fig.5D,E).

Next, we investigated if the higher purity of glycoconjugates that contained less unconjugated carrier protein, was able to trigger a RhaPS specific immune response. Three rabbits were immunised, and post-immunisation sera were collected at Day 35. Terminal sera were tested by ELISA for an anti-NanA-RhaPS response. All three post-immune bleeds were positive, whilst the three pre-immune bleeds did not detect the recombinantly produced RhaPS (SupplFig.6A). This was also confirmed by conducting ELISA analysis where plates coated with NanA-RhaPS antigen were recognised by the anti-NanA-RhaPS sera compared to pre-immune sera (SupplFig.6B). In order to determine whether the raised IgG antibodies can recognise the native antigen present on the surface of all Strep A bacteria, we conducted flow cytometry experiments with the immune rabbit sera. Various Strep A M-types and *S. dysgalactiae subsp. equisimilis* (SDSE) expressing GAC were assayed with sera from the NanA-RhaPS immunised rabbits and compared to the pre-immune sera. All tested Strep A strains showed binding by the anti-NanA-RhaPS immune sera. The number of cells detected with the antibodies as well as the flow cytometry histograms reveal that the NanA-RhaPS triggered antibodies recognise the tested strains, including M1_5448_ [6] the recently reported dominant M1_UK_ lineages [66], M75 serotype recently used in the human challenge study [67] and M89 (Fig.6). Overall, the results are in line with previous reports using chemically conjugated RhaPS backbone [6, 19, 20].

**Fig 6:**
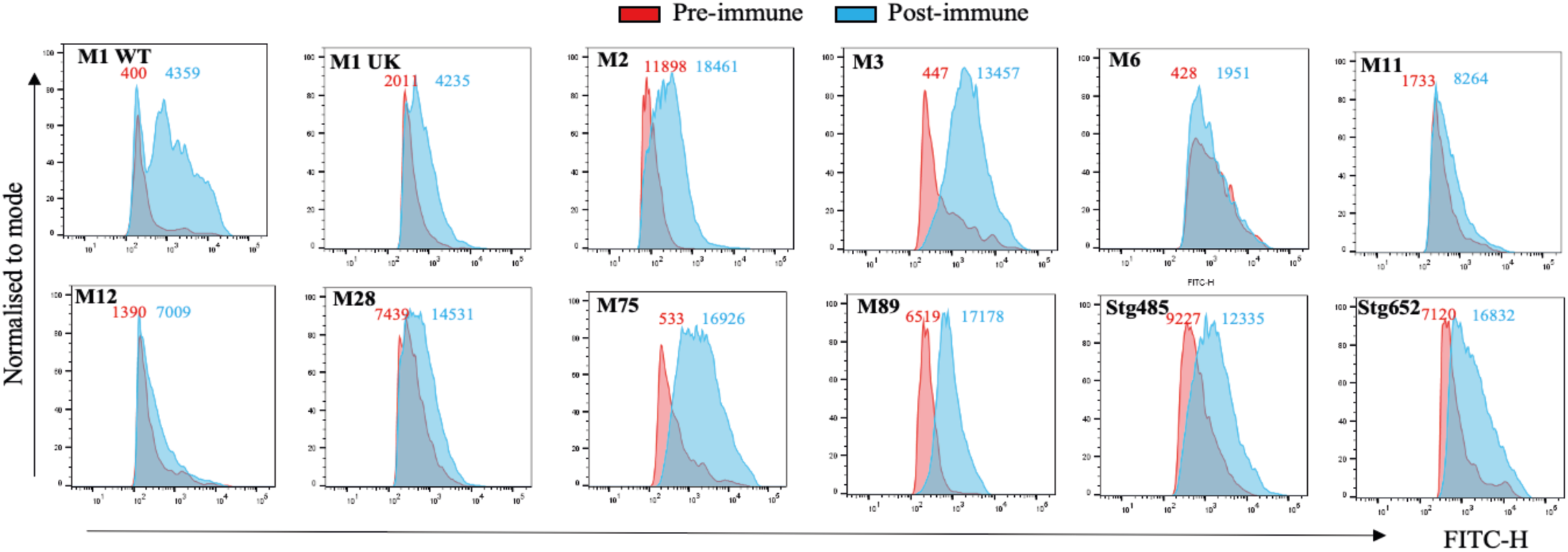
Representative histograms for antibody deposition on Strep A M serotypes and *S. dysgalactiae* subsp. *equisimilis* (SDSE) strain expressing GAC cluster (stg485 and stg652) were stained in rabbit antiserum (1:1000) for all strains except M2 (1:100) from pre-immune (red shading) and post-immune (NanA-RhaPS; blue shading) vaccinated groups. The number on the top shows the geometric mean fluorescence intensity (gMFI) of the pre-immune sera (red) and NanA-RhaPS immunised serum (blue) for the displayed histograms.

## DISCUSSION

There is currently no licensed vaccine for Strep A infection despite the importance of the disease globally and the reported increase in antimicrobial resistance. Bacterial polysaccharides are important vaccine antigens but to be effective they must be conjugated to a carrier protein. Such glycoconjugate vaccines demonstrate efficient antigen presentation and develop strong memory B cell with markedly high affinity antibody responses, which is especially important in young children where a Strep A vaccine is most sought.

The Lancefield Group A carbohydrate (GAC) is a conserved, abundant, and immunogenic surface structure that has been shown to be an excellent glycoconjugate vaccine candidate [6, 9, 19]. However, there is concern that the GlcNAc side chain may elicit unwanted immune complications [15]. Therefore, we have recombinantly produced the GAC backbone structure RhaPS that lacks the side chain and consists of the polyrhamnose homopolymer with the immunogenic →2)-α-L-Rha*p*-(1→3)-α-L-Rha*p*-(1→ disaccharide repeating unit, as a glycan component of potential Strep A vaccines [13]. We have previously shown that OMVs from *E. coli* that express RhaPS are strong antigens in the mouse infection model and generate specific bactericidal antibodies targeting all tested Strep A serotypes [9]. However, significant limitations of OMVs are that they can be heterologous and that they are difficult to manufacture. Similarly, Strep A glycoconjugate vaccines have been recently reported in conjugation of GAC to gold nanoparticles [68], an Ac-PADRE-lipid core [7] and a truncated Streptolysin O using click chemistry [18], but upscaling, non-natural amino acid incorporation and cost issues for production are major considerations for these approaches.

PGCT is an attractive low-cost technology platform to develop tailor-made glycoconjugate vaccines to combat Strep A but is untested due to the failure of current OTases to recognise the Strep A glycan structure. In this study, we have overcome the reducing end-group specificity limitation by engineering an OTase compatible disaccharide structures which enable the coupling of RhaPS to a selection of carrier proteins from different species. We engineered hybrid-glycan structures by combining the Strep A pathway with two characterised *Shigella* carbohydrate pathways. For the reducing end linkers employed herein *S. dysenteriae* type 1 has the structure α-L-Rha*p*-(1→3)-α-L-Rha*p*-(1→2)-α-D-Gal*p*-(1→3)-α-D-Glc*p*NAc, and *S. flexneri* 2a has the structure α-L-Rha*p*-(1→3)-α-L-Rha*p*-(1→3)-α-D-Glc*p*NAc. Much like GAC, glycan assembly takes place on the cytoplasmic side of the membrane on Und-PP. However, unlike GAC, the repeating units are flipped to the periplasm *via* the Wzx flippase[69] and are polymerised in the periplasm via the O-antigen polymerase, Wzy. We paired the respective enzymes with GacC-G to establish the immunodominant GAC repeating unit onto the respective *Shigella* reducing end groups.

These include a double-hit vaccine of the *S. pneumoniae* neuraminidase NanA-RhaPS, a dual-hit vaccine of inactivated protease IdeS-RhaPS, and the established toxoid carrier ExoA-RhaPS. We demonstrate that PGCT can be adapted to multiple carrier protein and glycan combinations, an important consideration in the development of glycoconjugate vaccines to avert the overuse of carrier proteins and avoid potential carrier-induced epitopic suppression.

Another novel feature of our study is the first synthesis of a peptidoglycan-anchored surface glycan (GAC/RhaPS) that is built to its entire length in the cytoplasm before being translocated via an ABC-transporter dependent pathway and therefore does not depend on repeating unit assembly in the periplasm prior to successfully being transferred to carrier proteins using PGCT [70]. This adds to the growing list of diverse bacterial glycans, such as capsular polysaccharides [37, 38], O-antigens [40] and glycoproteins [44] that can be used as the glycan component of PGCT-based glycoconjugate vaccines.

Immunisation studies with the recombinant NanA-RhaPS vaccine candidates trigger antibodies against the carrier protein and the RhaPS carbohydrate. We observed increased levels of cytokine mediators IFN-γ and IL-17 in NanA-RhaPS immunised mice over NanA immunsed animals (Fig. 5D,E). We suggest that these cytokine mediator increases in the glycoconjugate immunised animals are driven by carbohydrate-recognising T helper cells, suggesting the role of adaptive immune response against the RhaPS portion, providing valuable information on the ability of these recombinant glycoconjugate vaccine candidates to stimulate both protein and glycan immune responses. Similar observation and conclusions were made in a Group B Streptococcus type III glycoconjugate vaccine [21]. All tested *S. pyogenes* serotypes and the *S. dysgalactiae* subsp. *equisimilis* strains producing the Group A Carbohydrate are detected by antibodies generated against the recombinant NanA-RhaPS. Mice immunised with IdeS-RhaPS showed functional neutralising antibodies against IdeS in an IgG cleavage assay. The free protein content in these conjugate preparations may account for this functionality and further analysis with highly purified conjugates will determine whether there are truly functional carrier antibodies generated from IdeS/RhaPS. Sequential SEC purification was successful at markedly improving the purity of produced conjugates. However, further investigations should focus on improving the glycosylation efficiency within *E. coli* cells to streamline production and increase yields.

In conclusion, we demonstrate that glycoengineering leads to a novel opportunity to utilise PGCT to construct several promising Strep A glycoconjugate vaccines recombinantly. The coupling of RhaPS to diverse protein carriers confirms the success of the linker reducing end group strategy that could be more widely applied in the construction of other recombinant glycoconjugate vaccines. We currently also exploit related streptococcal pathogens that synthesise their respective surface carbohydrate *via* a related pathway containing a GacB homologous enzyme [13]. Our platform technology should allow the modification of these pathways analogous to this reported solution to modify the Strep A glycans and transform them into compatible substrates for the OTase enzymes and low-cost PGCT glycoconjugate production.

## METHODS

### Media and growth conditions

*E. coli* strains were cultured on LB agar and LB broth, Super Optimal broth (SOB: 20 g l^−1^ Tryptone, 5 g l^−1^ Yeast Extract, 2.4 g l^−1^ MgSO_4_, 0.186 g l^−1^ KCl, 0.5 g l^−1^ NaCl), Salty super Optimal broth (SSOB: 20 g l^−1^ Tryptone, 5 g l^−1^ Yeast Extract, 2.4 g l^−1^ MgSO_4_, 0.186 g/L KCl, 10 g/L NaCl), Brain Heart infusion (BHI) broth and 2x Yeast-Tryptone (16 g l^−1^ Tryptone, 10 g l^−1^ Yeast Extract, 5 g/L NaCl) at 37 °C or 28 °C as indicated. Antibiotics were added as necessary at the following concentrations: ampicillin 100 µg ml^−1^; erythromycin 150 µg ml^−1^; spectinomycin 40-80 µg ml^−1^; kanamycin 50 µg ml^−1^.

### Plasmid constructs

**Table 2.**

| Plasmid | Insert | Reference |
| --- | --- | --- |
| pHD0788 | pEXT21 (Empty plasmid) | [72] |
| pHD0890 | ExoA in pEXT21 | This study |
| pHD0769 | IdeS in pEXT21 | This study |
| pHD0256 | GAC-cluster | [13] |
| pHD0677 | Shi1 cluster | This study |
| pHD0768 | Shi2 cluster | This study |
| pHD0678 | Shi1 WbbP | This study |
| pHD0679 | Shi1 WbbPR | This study |
| pHD0680 | Shi1 WbbQR | This study |
| pHD0778 | NanA in pEXT21 | [37] |

### Cloning

Shi1 (pHD0677) and Shi2 (pHD0768), were subcloned from synthesized genes incorporated into cloning plasmids (IDT) and subcloned via the XbaI and NcoI sites into pBAD24 vector (pHD0131). IdeS (inactive) (pHD0769) was cloned from a synthetic gene block and inserted into XbaI and EcoRI linearised empty plasmid pEXT21 (pHD0788) via restriction less cloning (InFusion, Takara). A signal-peptide was included 5’ of the IdeS fragment. The Shi1 derivative plasmids pHD0678-680 were obtained via PCR and NcoI and XbaI digestion using the primers listed and the pHD0677 as DNA template. All constructs were verified by DNA Sanger sequencing using the DSTT University of Dundee sequencing facility and/or via whole plasmid sequencing (Plasmidsaurus). ExoA was cloned into the pEXT21 vector via the EcoRI and BamHI sites with primers as shown below.

#### Primers

pHD0678_679_fwd CCATAGCATTTTTATCCATAAGATTAGCG

WbbP_XbaI_Rev CGACTCTAGATTAATCAGGAATCCCTAGTATTTTTAATAACTTTAC

WbbR_XbaI_rev CGACTCTAGATTACATTTTAAGACCATCTTTTATTCCTTTTAAATATAAATGC

pHD0680_fwd ATTCCATGGATGAACAAGTACTGCATCCTTGTGC

pHD0680_rev CATGGGGTCAGGTGGGAC

pHD0890_fwd CTAGCGCCGCCGATCAG

pHD0890_rev GAGGATCCTCAGTGGTGGTGG

### Small scale glycoconjugates expression and purification

Glycoconjugate test expressions were conducted in rfaS cells [13]. Transformed cells were grown at 37°C overnight in LB media, induced with 0.1 mM IPTG and 0.2% arabinose. *E. coli* MAGIC cells containing a chromosomal copy of the *pglB* gene under an IPTG inducible promotor (*cedA*::*pglB*) [53, 54] were made electro- or chemically competent in LB media containing Kanamycin (50 µg/mL) and transformed with 10-100 ng of plasmids of interest (Table 1) and selected on LB agar supplemented with the corresponding selection antibiotics Kanamycin (K, 50 µg/mL), Ampicillin (A, 100 µg/mL), Spectinomycin (S, 80 µg/mL) and Erythromycin (E, 150 µg/mL), KASE. One to five colonies were swabbed of the plate and grown overnight in LB media with the required antibiotics at 37°C. Next morning, they were diluted 1:100 in fresh 10 ml of the specified growth medium supplemented with 0.2% (w/v) L-arabinose (Sigma) and 1 mM glycerol (Sigma) in 30 ml closed tubes. Cultures were grown at 28°C, 180 rpm to OD_600_ 0.5 at which point expression of inducible proteins was promoted by the addition 1 mM IPTG (for pEXT21 vectors and chromosomally integrated *pglB*). Cultures were then grown at 28°C for further 16 h followed by cell harvesting by centrifugation at 5300 x *g*, 4°C for 15 min. For His-purifications pellets were resuspended in 1 ml lysis buffer (50 mM NaH_2_PO_4_, 300 mM NaCl, and 10 mM imidazole, pH 8.0). Cells were lysed using BeadBug zirconium lysing tubes (Sigma) in a FastPrep homogeniser (MP Biomedicals) and insoluble material was removed by pelleting. Histidine tagged proteins were subsequently purified from cell lysates by Ni-NTA affinity chromatography (resin) and eluted in 50 mM Tris HCl pH 8, 300 mM NaCl, 300 mM imidazole. Glycoconjugate production was verified by SDS-PAGE followed by western blotting and ELISA as described below.

### Large scale glycoconjugate production and purification

The production of IdeS-RhaPS and NanA-RhaPS was upscaled for downstream analysis including immunisation studies, NMR, and mass spectrometry. The strains were grown in LB + KASE media overnight at 37°C and back diluted 1:100 into prewarmed SOB or THY media containing the same antibiotics. At OD600 ∼ 0.6, the cells were shifted to 28°C and induced with an induction cocktail containing 0.1 mM IPTG, 0.2% arabinose, 1 mM GlcNAc and 10% glycerol. After 16-18 h at 28°C, the cultures were harvested by centrifugation at 4200 rpm, 30 min, 4°C and pellets resuspended in PBS buffer supplemented with protease inhibitors, lysozyme, 10 mM Imidazole and 2 mM TCEP. Cells were lysed using a constant cell disrupter. After three passages at 30 kpsi, the samples were subjected to centrifugation at 30k x *g*, 30 min, 4°C and the supernatant was bound to 5 mL HP IMAC/NTA columns, charged with Co^2+^. The bound protein was washed with PBS buffer containing 15 mM Imidazole and 0.2 mM TCEP and eluted with PBS buffer containing 500 mM Imidazole and 0.2 mM TCEP. Fractions eluted from the NTA column were checked by SDS-PAGE and those that contained glycoconjugates were concentrated by centrifugation with a 10 kDa MWCO concentrator and injected into Superdex 75 16/600 column equilibrated in endotoxin free PBS. Peak fractions were analysed by SDS-PAGE and concentrated with a 10 kDa MWCO concentrator to 5 mg/mL. The mixture of free carrier protein and glycoconjugate was pooled and concentrated for mice immunisation studies.

Samples for rabbit immunisation studies were processed *via* an additional Superdex 75 16/600 purification step. Peak fractions were pooled that only contained glycosylated proteins and not the free carrier protein alone as judged by SDS-PAGE (Coomassie stain). Protein samples were flash frozen in 25% glycerol until immunisation.

Salts were removed for NMR analysis via high-performance liquid chromatography (HPLC). Glycoconjugate aliquots of 75 µL to 5 mL (3.5-5 mg/mL in 25% glycerol) were injected and passed over a C4 column (either Perkin Elmer Epic C4 column or alternatively Phenomenex Jupiter 10u C4 300Å 250−10 mm 10 micron). A solvent gradient was applied using solvent A (H_2_O plus 0.1% (v/v) TFA), solvent B (acetonitrile 0.1% (v/v) TFA) at 3 mL/min with start condition of 95%A and 5% B. The linear gradient was gradually increased to 90% B. Fractions containing the protein samples were identified via OD220 nm, confirmed by SDS-PAGE analysis, pooled, and freeze dried.

### Glyco-conjugate analysis *via* SDS-PAGE and Western blotting

His-pulldowns and OD_600_-matched periplasmic extracts were mixed with LDS sample buffer and denatured at 95°C for 10 min before being separated on 4-12 % bis-tris gels in MOPS or MES buffer (Invitrogen, USA). Gels were then electroblotted onto a nitrocellulose membrane, which was then washed in PBS with 0.1% Tween-20 (PBS-T) and incubated overnight at 4°C shaking in PBS-T with rabbit anti-GAC antibodies (Abcam ab21034) at a 1:5,000 dilution and mouse anti-His monoclonal antibody (Thermo Fisher Scientific or Abcam ab216773, Goat pAb to rabbit IgG)) at a 1:5,000 dilution to detect RhaPS and His-tagged carrier proteins. Alternatively, His-tagged proteins were detected by ab117504 (mouse mAb to His-tag). Blots were washed three times in PBS-T, followed by incubation for 30 min at room temperature with goat anti-rabbit IRDye 800CW and goat anti-mouse IgG IRDye 680 (LI-COR Biosciences) at 1:10,000 dilution. After three final washes, fluorescent signals in two channels, 700 and 800 nm, were detected with an Odyssey CLx LI-COR detection system (LI-COR Biosciences).

### Dot-blotting

*E. coli* cells were grown in 5 ml cultures of the specified growth medium supplemented with appropriate antibiotic and 0.2% (w/v) L-arabinose shaking at 180 rpm at 37 °C overnight. Cultures were normalised to OD600 nm = 5 and were sedimented by centrifugation (6,000 x *g*, 10 minutes) and pellets were washed three times with sterile PBS. Cell suspensions (2.5 µl) were spotted onto nitrocellulose membrane and air dried for 30 minutes. Membranes were washed with PBST and probed with anti-GAC antibody (abcam9191) at a 1: 5,000 dilution for 45 minutes. Membranes were washed three times with PBST and incubated with goat anti-rabbit IRDye 800CW (LI-COR Biosciences) antibody at a dilution of 1 in 10,000 in PBST for 30 min. Fluorescent signal was detected with the Odyssey LI-COR detection system (LI-COR Biosciences).

### NMR spectroscopy

The NMR spectra were recorded on a Bruker AVANCE III 700 MHz spectrometer equipped with a 5 mm TCI Z-Gradient Cryoprobe (^1^H/^13^C/^15^N). A sample of the glycoconjugate construct IdeS-RhaPS (10 mg) was dissolved in D_2_O (0.6 mL) and analysed using a 5 mm o.d. NMR tube at 298 K. Chemical shifts are reported in ppm using external sodium 3-trimethylsilyl-(2,2,3,3-^2^H_4_)-propanoate in D_2_O (TSP, *δ*_H_ 0.00 ppm) and external 1,4-dioxane in D_2_O (*δ*_C_ 67.40 ppm) as references. 2D NMR experiments, viz., ^1^H,^13^C-HSQC[73] and ^1^H,^13^C-HMBC[74] with a delay for the evolution of the long-range couplings corresponding to 12.5 Hz were performed with standard Bruker pulse sequences using TopSpin 3.2 software for acquisition of data and TopSpin 4.1.3 for processing and analysis of data.

### Mass spectrometry

The purified NanA-RhaPS and IdeS-RhaPS proteins were concentrated to 3.1 mg/mL and 3.5 mg/mL, respectively. Thirty micrograms of individual glycoprotein standards were resuspended in 200 mM HEPES pH 8, 40 mM Tris(2-carboxyethyl)phosphine Hydrochloride (TCEP), 40 mM chloroacetamide (CAA) for reduction and alkylation of cysteine residues at 95°C for 5 min. Prior to trypsin digestion, the samples were subjected to acetone precipitation. The protein pellet was resolubilized in 50 mM TEAB buffer, and trypsin was added in a 1:50 ratio (enzyme:substrate). After overnight incubation at 37°C, the resulting glycopeptide/peptide mixtures were dried in the speed vac without additional heating. In the case of chymotrypsin, the samples were incubated at 25°C for 4 h.

### Reverse Phase LC-MS/MS

All samples were resuspended in water containing 0.1% Formic acid. Samples were separated by RP-HPLC using a Thermo Scientific Dionex UltiMate 3000 system coupled to Thermo Scientific™ Orbitrap Exploris™ 480 equipped with a FAIMS Pro device (Thermo Fisher Scientific) with Instrument Control Software version 3.3.

Samples were loaded onto the PepMap C-18 trap-column (0.075 mm × 50 mm, 3 μm particle size, 100Å pore size (Thermo Fischer Scientific)) 20 μL for 5 minutes with buffer A (0.1% formic acid, 2% acetonitrile) and separated over a 15 cm × 75 μm C18 column (column material: ReproSil C18,1.9 μm (Pepsep)) at 300 nL/min flow rate. For the nLC-MS/MS methodsology the duration of each experiment wes 60 min total. The separation of peptides and modified peptides were achieved by altering the buffer composition from 3% buffer B (0.1% formic acid, 80% acetonitrile) to 30% B over 23 min, then from 30% B to 45% B over 3 min, and then a rapid increase from 45% B to 98% B over 30 sec. The composition was held at 98% B for 3 min and then reduced to 2% B over 30 sec before being held at 3% B for another 20.5 min.

All data were collected in positive ion mode using Data Dependent Acquisition mode. FAIMS separations were performed using the following settings: inner and outer electrode temperature = 100°C (except where noted); Static FAIMS analysis was carried out using CVs of −20, −30, −35, −40, and −50. In all cases, 400 ng peptide was injected per analysis.

The ion source was held at 2200 V compared to the ground, and the temperature of the ion transfer tube was held at 275°C. Survey scans of precursor ions (MS1) were collected in the mass-to-charge (*m/z*) range of 350 – 2,000 at 60,000 resolution with a normalized AGC target of 250% and a maximum injection time of 60 ms, RF lens at 50%, data acquired in profile mode. Monoisotopic precursor selection (MIPS) was enabled for peptide isotopic distributions, precursors of z = 2-8 were selected for data-dependent MS/MS scans for 3 seconds of cycle time (unless noted otherwise) with an intensity threshold of 4.0e3, and dynamic exclusion was set to 40 seconds with a ±10 ppm window set around the precursor monoisotopic. Also, Precursor Fit was enabled for the MS/MS scans at the threshold of 70% within the 2 Da window.

The mass spectrometry proteomics data have been deposited to the ProteomeXchange Consortium via the PRIDE[75] partner repository with the dataset identifier PXD037039.

### Data Analysis - Peptide and Glycopeptide Identification

Raw data files were batch-processed using Byonic v4.3.4 (Protein Metrics Inc. ([76])) with the proteome databases, as given in SupplTable1.

For all searches, a semitryptic (slow) specificity was set, and a maximum of two missed cleavage events were allowed. Carbamidomethyl was set as a fixed modification of cysteine, while oxidation of methionine and Protein N-terminal acetylation was included as a variable modification. A maximum mass precursor tolerance of 10 ppm was allowed, while a mass tolerance of up to 20 ppm was set for HCD fragments. Wildcard search was enabled for all peptides in the mass range of 100 to 10,000 Da restricted to asparagine. Searches were performed with either Trypsin or Chymotrypsin cleavage specificity towards each protein sample, with a maximum false discovery rate (FDR) of 1.0% set for protein identifications.

The mass spectrometry proteomics data have been deposited to the ProteomeXchange Consortium via the PRIDE partner repository with the dataset identifier PXD037039. Identified modified peptides were inspected manually, and only peptides following strict protease cleavage specificity were considered for validation and quantitation. For site-specific relative quantification, the extracted ion chromatogram of the identified peaks was calculated using Skyline[77]. Each peptide that contains PTM sites was normalized individually so that the sum of all its proteoform areas was set at 100%.

Peptide sequence coverage was verified using FragPipe computational platform (v18)[78] with MSFragger (v3.5), Philosopher (v4.4.0). Peptide identification was performed using MSFragger search engine using .raw files. The proteome database listed in SupplTable2 was used for identification. Reversed and contaminant protein sequences were appended to the original databases as decoys using the default function. Enzyme specificity was set to either ‘stricttrypsin’ or ‘Chymotrypsin’. Up to two missed cleavages were allowed. Isotope error was set to 0/1/2/3. Peptide length was set from 7 to 50, and peptide mass was set from 500 to 5000 Da. Oxidation of methionine, and acetylation of protein N-termini was set as variable modifications. Carbamidomethylation of Cysteine was set as a fixed modification. A maximum number of variable modifications per peptide was set to 3.

### Monosaccharide composition analysis by GC-MS

An amount of 0.56 µg of IdeS-RhaPS glycoconjugate was mixed with 2 nmoles of *scyllo*-Inositol as an internal standard, split into three aliquots and subjected to methanolysis, trimethylsilylation, and monosaccharide composition analysis, as described earlier [79]. The TMS derivatized samples were analyzed by GC-MS (Agilent Technologies, 7890B Gas Chromatography system with 5977A MSD, equipped with Agilent HP-5ms GC Column, 30 m × 0.25 mm, d_f_ 0.25 µm).

### Ethical approval

Mice studies were approved by the University of Dundee welfare and ethical use of animals committee and conducted in accordance with the UK Home Office approved project licence [PPL PEA2606D2]. The experiments involving rabbits were conducted in association with Davids Biotechnologie GmbH, Germany.

### Murine and rabbit immunisation studies

Mouse Model: Five- to six-week-old female C57BL/6J mice were acquired from Charles River Laboratories, UK. The animals were acclimatised for 10 days prior to immunisation.

Mice were immunised subcutaneously either with NanA-RhaPS (n=10) or NanA (n=10) or IdeS-RhaPS (n=10). Identical booster injections were administered on day 14 and day 28. Animals were euthanised by CO_2_ asphyxiation and bled by cardiac puncture on day 70. Placebo vaccinated mice (n=10) were used for baseline measurements. All groups received Sigma adjuvant system at 1:1 ratio to immunogens.

Rabbit Model: Three New Zealand White rabbits were immunised with three equal doses of NanA-RhaPS (10 µL of 3.5 mg/mL diluted in succinate buffer adjuvanted with Adju-phos® (Invivogen)). Immunizations (250 µL per injection per dose) were performed on day 0, 14, and 28 with terminal bleed for serum performed on day 35. PBS/injection) on day 0. The animals were culled by CO_2_ asphyxiation and bled by cardiac puncture. All immunisation work was conducted by Davids Biotechnologie GmbH.

### ELISA

Polysaccharide specific IgG were measured by coating the Nunc Maxisorp 96 well plate with 1 µg/well with either purified NanA-RhaPS or IdeS-RhaPS recombinant protein in endotoxin free PBS and the plates were incubated overnight at 4 °C. Next day, the plates were washed before adding the murine or rabbit antiserum 50 µl/well at 1:1,000 dilution and incubated for an hour at room temperature. The unbound antibodies were removed by washing the plates with ELISA wash buffer (PBS with 0.05% Tween 20). Bound antibodies were probed using 50 μl/well of 1:1,000 of anti-mouse IgG HRP (Sigma–Aldrich) and incubated for an hour at room temperature; 75 μl/well of tetramethylbenzidine substrate (Sigma–Aldrich) were used for HRP detection and the reaction was stopped using 75 μl/well of 1 M H_2_SO_4_, and absorbance read at 450 nm.

### Immunoblot analysis using rabbit sera

Antisera from NanA-RhaPS immunised rabbits were tested for binding to *E. coli* cells or Strep A by spot blot and western blot analysis. Overnight cultures of *E. coli* cells producing the *Streptococcus mutans* and Strep A RhaPS from their respective gene clusters [13] on their outer membrane were spotted onto nitrocellulose membrane, dried and blocked with 5% non-fat dried milk in Tris-Buffered Saline, 0.1% Tween® 20 (TBS-T). Rabbit serum was diluted 1:500 in TBS-T. The membranes were incubated with the rabbit sera for 1 h at room temperature. The membranes were then washed three times with TBS-T and the fluorescent labelled IRDye anti-rabbit secondary antibody was used for both blots, ab216773 (Abcam) in a 1:5000 dilution. After 1 h incubation, the membranes were washed as above and imaged using a LICOR instrument.

### Flow cytometry

Antibody binding to the GAS surface were conducted using flow cytometry analysis as previously described in [51]. Briefly, overnight Strep A cells or *S. dysgalactiae* subsp. *equisimilis* (SDSE) isolates expressing Group G Carbohydrate (SDSE_GGC) gene cluster or SDSE isolates which have replaced the GGC gene cluster with a functional Group A Carbohydrate cluster (SDSE_GAC) were diluted 1:10 the next day. The cells were washed with 0.3% BSA in PBS (flow buffer) at OD_600nm_ −0.4. Non-specific binding sites on the bacterial cells were blocked using human IgG (Sigma, UK) for an hour and washed with flow buffer again before incubated overnight with the rabbit immune sera (1:1,000 dilution). Following washes, the cells were resuspended in the detection antibody (Alexa fluor 488-conjugated goat anti-Rabbit IgG; Thermo Fisher, UK); 4% PFA was used to fix the cells before analysing by flow cytometry (BD Bioscience). A total of 20,000 events were acquired and the histograms were analysed using FlowJo software version 10.6.2.

### IdeS Neutralisation Assay

An ELISA method was adapted from a previously published method[64]. Maxisorp 96 well ELISA plates (Nunc) were coated with 1/20,000 dilution of Goat anti-Human Fab (Sigma) overnight at 4°C. 10 ng recombinant IdeS was pre-incubated with sera dilutions as indicated at 37°C 1 h before addition of 100 ng human IgG (Sigma) in assay diluent (AD: 1% BSA PBS-T 0.3% Tween-20) and further incubation at 37°C for 3 h. ELISA plates were blocked for 1 h in AD before addition of assay samples for 1 h followed by incubation with 1/20,000 dilution Goat anti-human Fc-HRP (Sigma). All steps were conducted at 37°C and separated by three washes with PBS-T (0.05% Tween-20). Plates were detected with 100 μl TMB (3,3’,5,5’-tetramethylbenzidine, ThermoFisher Scientific) and incubated at RT for 15 min and the reaction was stopped with 100 μl of 1 M sulphuric acid. Plates were read at 450 nm on a SoftMax Pro plate reader. Signal was compared with no sera assay conditions, non-immune normal mouse sera (NMS) and a positive control serum.

### Cytokine assays

Determination of cytokine mediators from the mouse splenocytes stimulated with the glycoconjugate vaccine were conducted following the protocol according to Moffit *et al.*, [80]. Briefly, freshly isolated mice spleen was flushed using RPMI media containing glutamine (Gibco), 10% FBS (Gibco), and 100 U/ml Penicillin-Streptomycin (ThermoFisher). The harvested cells were passed through a 40 µm cell sieve and washed further with 5 ml of RPMI media by spinning the cells at 300 x g for 10 minutes. To the pellet, 1 ml of RBS lysis buffer (BioLegend) was added and then incubated for 1-2 minutes. To the cells, 10 ml of RPMI media was added and spun down at 300 x g for 10 minutes. The resulting pellet was resuspended in 1 ml of 1 ml of RPMI media, enumerated, and adjusted to 7.5 x 10^6^ cells/ml. 200 µl of cells (1 x 10^6^ cells/ml) were added to the designated wells in the 96 well plate and 50 µl of 50 µg/ml of recombinant protein (NanA-RhaPS) was added and left to incubate at 37°C 5% CO_2_ for 3 days. The cells were pelleted (400 x g for 10 minutes) and used further for analysis of cytokine mediators IFN-γ and IL-17 (ThermoFisher) for ELISA analysis. All steps were conducted at room temperature.

## Data Availability

The mass spectrometry proteomics data have been deposited to the ProteomeXchange Consortium via the PRIDE partner repository [http://proteomecentral.proteomexchange.org/cgi/GetDataset] with the dataset identifier PXD037039.

## Acknowledgements

We thank Dr Emily Kay (LSHTM) for glycoengineering technical advice.

## Author Contributions

S.A.C, I.P., D.N., H.A.S., A.R., K.B., S.T., R.N., K.A., K.L., S.A., M.R., U.S-L., G.W., H.C.D. performed the experiments and collected and analysed the data. F.M., G.W., B.W.W. analysed and interpreted data. B.W.W, H.C.D, I.P. and S.A.C wrote the manuscript. B.W.W and H.C.D supervised the study. All authors read and approved the final manuscript.

