## Supplementary Figures for "A platform for the recombinant production of Group A Streptococcus glycoconjugate vaccines"

Supplementary Figure 1)

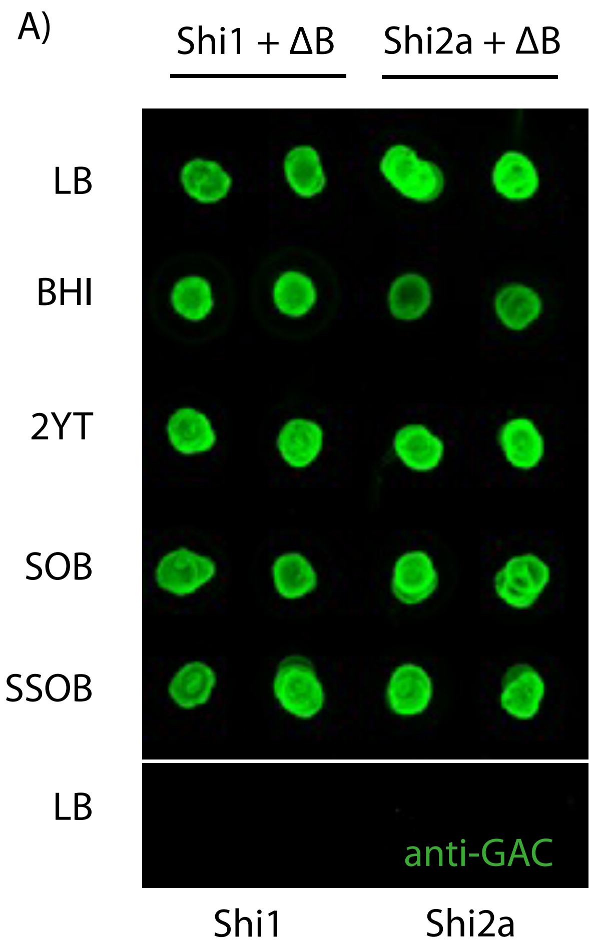


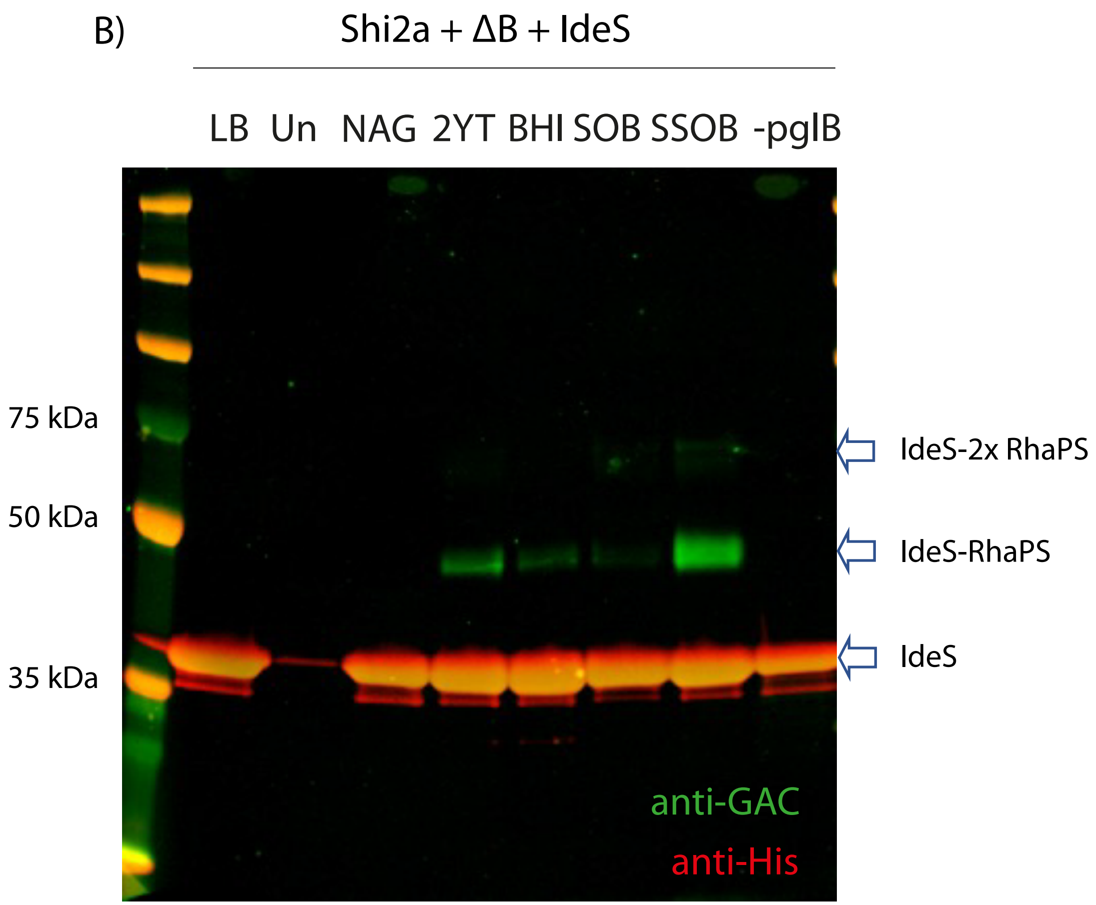

Supplementary Figure 2)

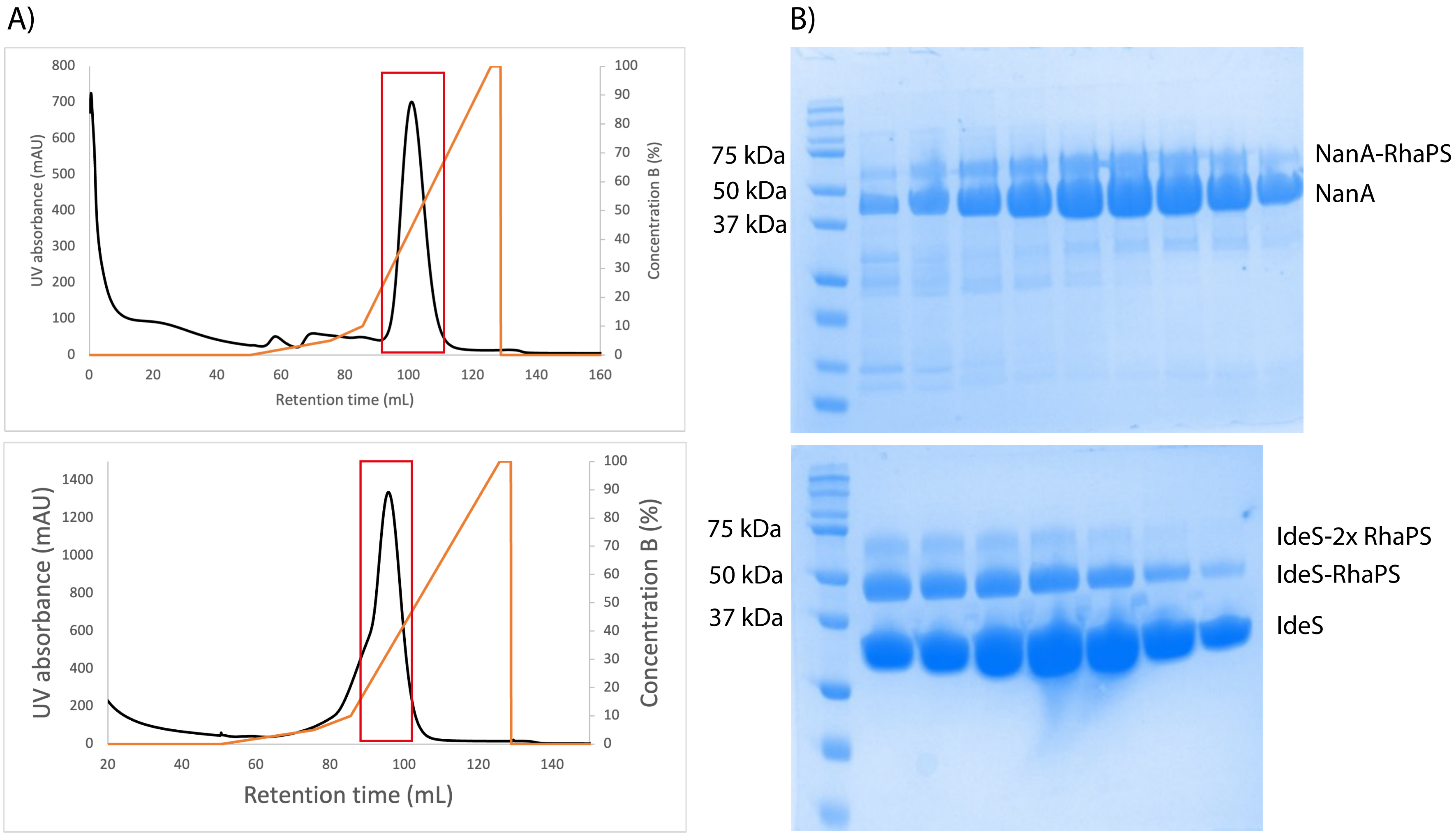

Supplementary Figure 3)

A)

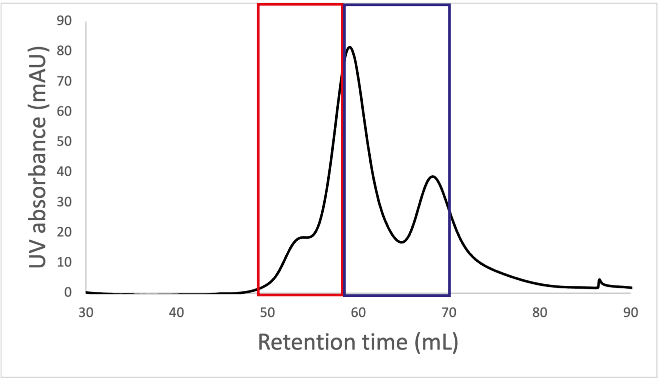


B)

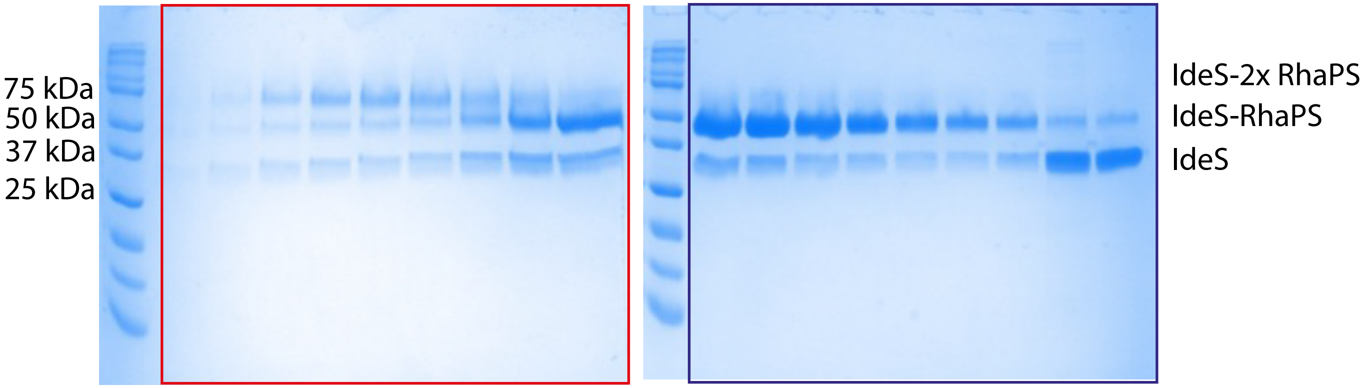

Supplementary Figure 4)

A)

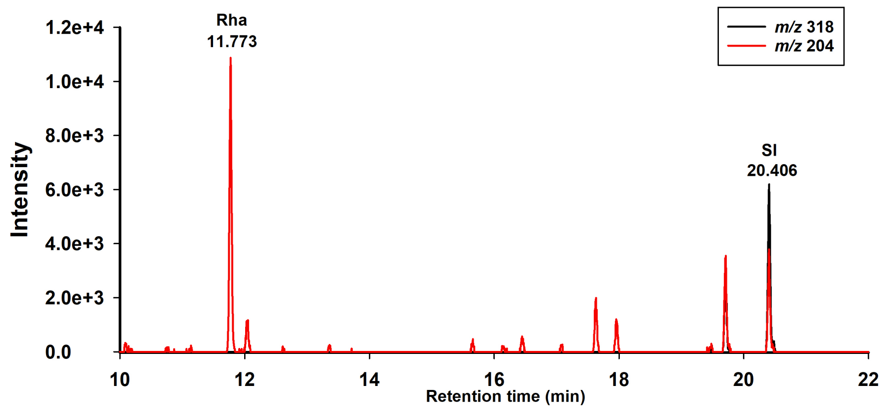


B)

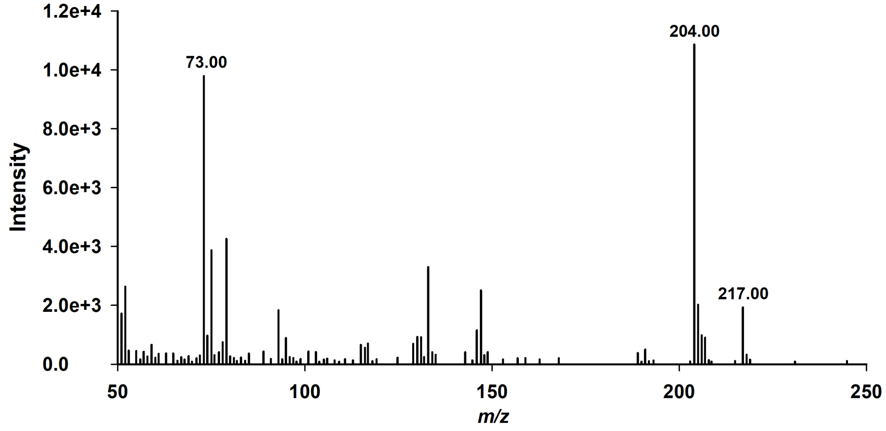

### Supplementary Figure 5)

A)

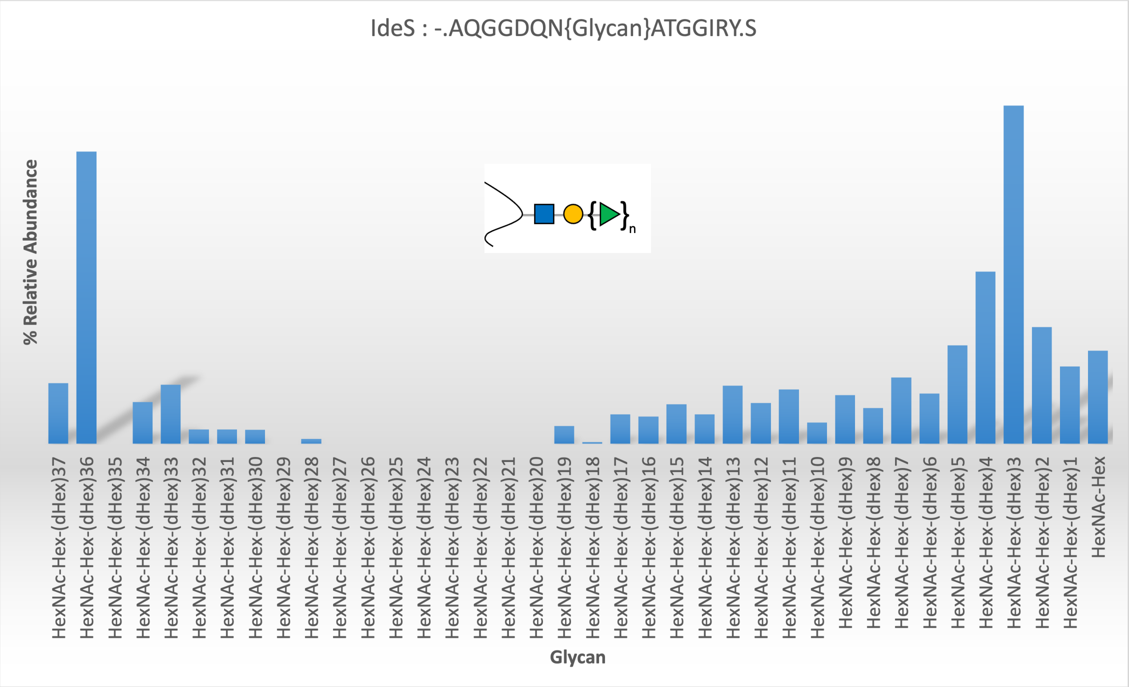


B)

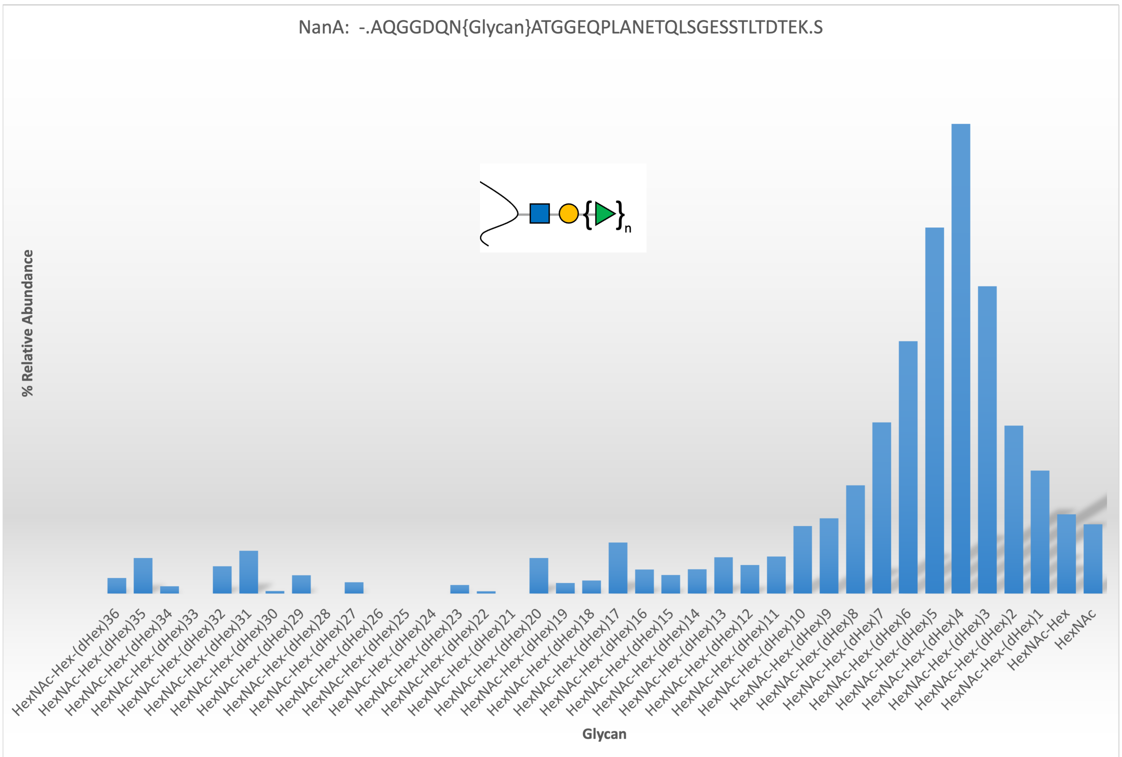

Supplementary Figure 6)

A)

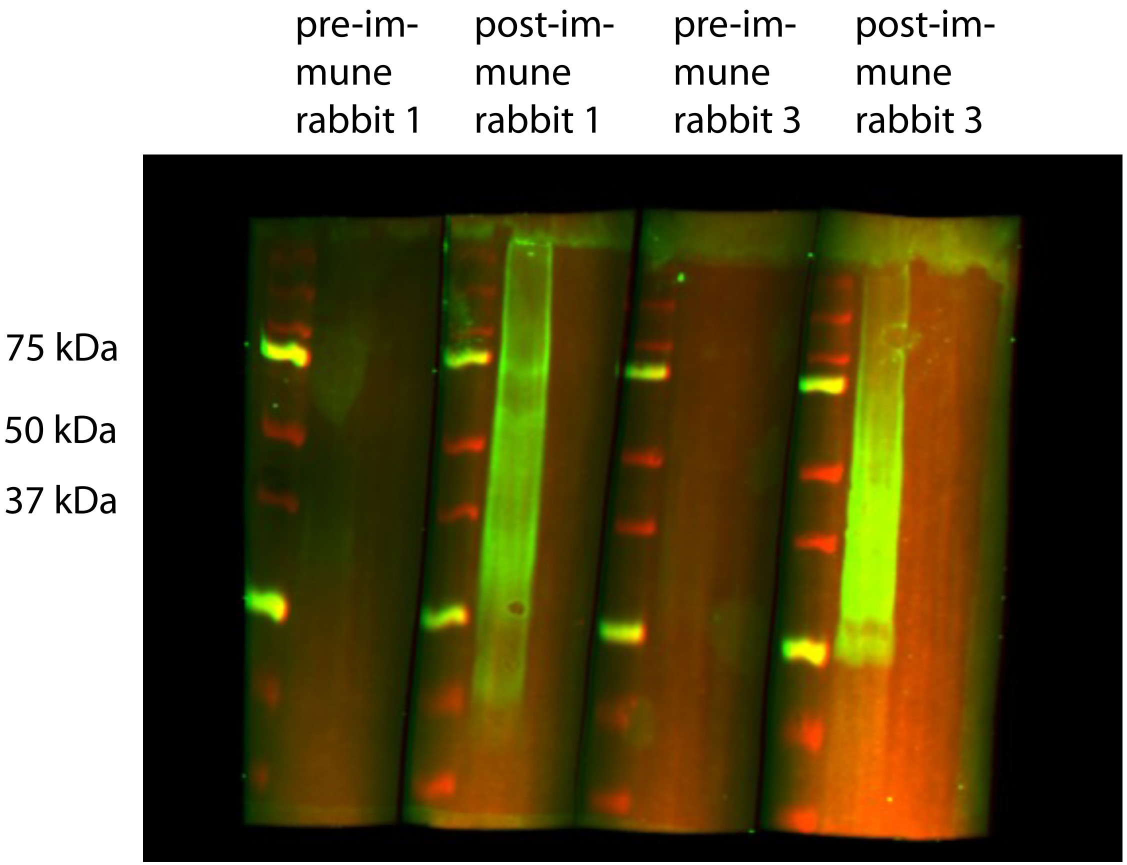


B)

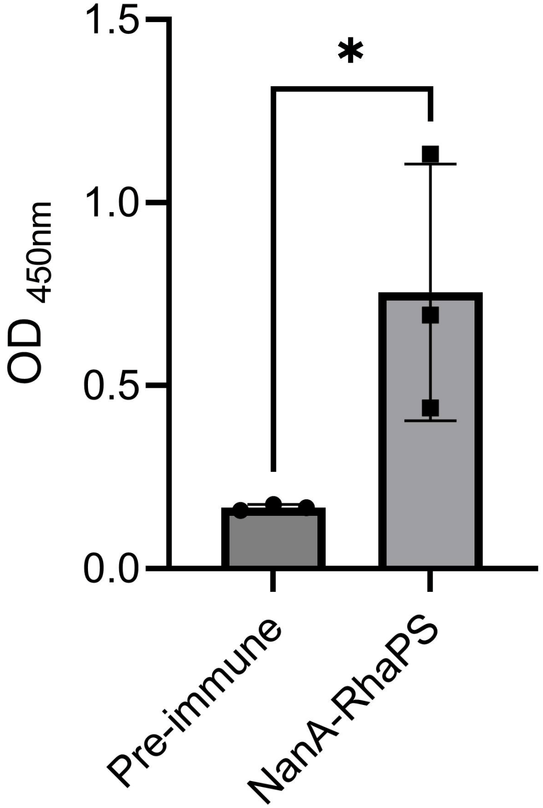
