## Supplemental Table for "A platform for the recombinant production of Group A Streptococcus glycoconjugate vaccines"

### GP-rhamnan

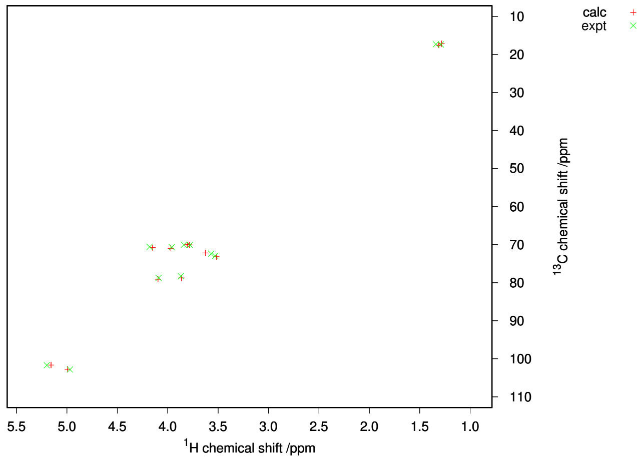

#### Predicted $^{13}$C and $^{1}$H NMR chemical shifts

##### Structure

→2) α-L-Rha$^{ii}$ (1→3) α-L-Rha$^{i}$ (1→

| | 1 | 2 | 3 | 4 | 5 | 6 |
|---|---|---|---|---|---|---|
| →3) α-L-Rha$^{i}$ (1→ | 1 | 2 | 3 | 4 | 5 | 6 |
| Expected Calc. Error: 3.06 | 102.76 | 70.81 | 78.75 | 72.19 | 70.05 | 17.20 |
| | 4.99 | 4.15 | 3.86 | 3.63 | 3.79 | 1.28 |
| →2) α-L-Rha$^{ii}$ (1→ | 1 | 2 | 3 | 4 | 5 | 6 |
| Expected Calc. Error: 2.53 | 101.66 | 79.09 | 70.97 | 73.18 | 70.02 | 17.57 |
| | 5.16 | 4.10 | 3.97 | 3.52 | 3.81 | 1.31 |

#### Assignment of $^{13}$C, $^{1}$H resonances

| Experimental | Predicted | Expt-Pred | Assignment |
|---|---|---|---|
| 102.82 - 4.97 | 102.76 - 4.99 | 0.03 | α-L-Rha$^{i}$ - 1 |
| 101.70 - 5.20 | 101.66 - 5.16 | 0.04 | α-L-Rha$^{ii}$ - 1 |
| 78.76 - 4.09 | 79.09 - 4.10 | 0.07 | α-L-Rha$^{ii}$ - 2 |
| 78.31 - 3.87 | 78.75 - 3.86 | 0.09 | α-L-Rha$^{i}$ - 3 |
| 72.94 - 3.53 | 73.18 - 3.52 | 0.05 | α-L-Rha$^{ii}$ - 4 |
| 72.40 - 3.57 | 72.19 - 3.63 | 0.07 | α-L-Rha$^{i}$ - 4 |
| 70.67 - 3.96 | 70.97 - 3.97 | 0.06 | α-L-Rha$^{ii}$ - 3 |
| 70.61 - 4.18 | 70.81 - 4.15 | 0.05 | α-L-Rha$^{i}$ - 2 |
| 70.10 - 3.78 | 70.05 - 3.79 | 0.01 | α-L-Rha$^{i}$ - 5 |
| 70.04 - 3.84 | 70.02 - 3.81 | 0.03 | α-L-Rha$^{ii}$ - 5 |
| 17.35 - 1.34 | 17.57 - 1.31 | 0.05 | α-L-Rha$^{ii}$ - 6 |
| 17.38 - 1.29 | 17.20 - 1.28 | 0.04 | α-L-Rha$^{i}$ - 6 |
| Error=0.59 (0.05/signal) , RMS error=0.05. | | | |

#### Assignment of long range $^{13}$C, $^{1}$H correlations

| Experimental | Predicted | Expt-Pred | Assignment |
|---|---|---|---|
| 102.82 - 4.09 | 102.76 - 4.10 | 0.01 | α-L-Rha$^{i}$ - 1, α-L-Rha$^{ii}$ - 2 |
| 101.70 - 3.87 | 101.66 - 3.86 | 0.01 | α-L-Rha$^{ii}$ - 1, α-L-Rha$^{i}$ - 3 |
| 78.76 - 4.97 | 79.09 - 4.99 | 0.07 | α-L-Rha$^{ii}$ - 2, α-L-Rha$^{i}$ - 1 |
| 78.31 - 5.20 | 78.75 - 5.16 | 0.10 | α-L-Rha$^{i}$ - 3, α-L-Rha$^{ii}$ - 1 |
| 78.31 - 4.97 | 78.75 - 4.99 | 0.09 | α-L-Rha$^{i}$ - 3, α-L-Rha$^{i}$ - 1 |
| 70.67 - 5.20 | 70.97 - 5.16 | 0.07 | α-L-Rha$^{ii}$ - 3, α-L-Rha$^{ii}$ - 1 |
| 70.10 - 4.97 | 70.05 - 4.99 | 0.03 | α-L-Rha$^{i}$ - 5, α-L-Rha$^{i}$ - 1 |
| 70.04 - 5.20 | 70.02 - 5.16 | 0.04 | α-L-Rha$^{ii}$ - 5, α-L-Rha$^{ii}$ - 1 |
| Error=0.42 (0.05/signal) , RMS error=0.04. | | | |

#### Assigned experimental $^{13}$C and $^{1}$H NMR chemical shifts

##### Structure

→2) α-L-Rha$^{ii}$ (1→3) α-L-Rha$^{i}$ (1→

| | 1 | 2 | 3 | 4 | 5 | 6 |
|---|---|---|---|---|---|---|
| →3) α-L-Rha$^{i}$ (1→ | 1 | 2 | 3 | 4 | 5 | 6 |
| $^{13}$C Error: 1.14. $^{1}$H Error: 0.13. | 102.82 | 70.61 | 78.31 | 72.40 | 70.10 | 17.38 |
| | 4.97 | 4.18 | 3.87 | 3.57 | 3.78 | 1.29 |
| →2) α-L-Rha$^{ii}$ (1→ | 1 | 2 | 3 | 4 | 5 | 6 |
| $^{13}$C Error: 1.15. $^{1}$H Error: 0.13. | 101.70 | 78.76 | 70.67 | 72.94 | 70.04 | 17.35 |
| | 5.20 | 4.09 | 3.96 | 3.53 | 3.84 | 1.34 |

Generated 2022-07-11 19:51:49+02:00.

NMR chemical shift data of GP-rhamnan

| HSQC | | HMBC | |
|---|---|---|---|
| 1H [ppm] | 13C [ppm] | 1H [ppm] | 13C [ppm] |
| 1.34 | 17.35 | 5.20 | 70.04 |
| 1.29 | 17.38 | 5.20 | 70.67 |
| 3.84 | 70.04 | 5.20 | 78.31 |
| 3.78 | 70.10 | 4.97 | 70.10 |
| 4.18 | 70.61 | 4.97 | 78.31 |
| 3.96 | 70.67 | 4.97 | 78.76 |
| 3.57 | 72.40 | 4.09 | 102.82 |
| 3.53 | 72.94 | 3.87 | 101.70 |
| 3.87 | 78.31 | | |
| 4.09 | 78.76 | | |
| 5.20 | 101.70 | | |
| 4.97 | 102.82 | | |
| | | 1.34 | 70.04 |
| | | 1.34 | 72.94 |
| | | 1.29 | 70.10 |
| | | 1.29 | 72.40 |
